# Computational modelling of cell motility modes emerging from cell-matrix adhesion dynamics

**DOI:** 10.1101/2021.06.09.447692

**Authors:** Leonie van Steijn, Clément Sire, Loïc Dupré, Guy Theraulaz, Roeland M.H. Merks

**Affiliations:** Mathematical Institute, Leiden University, Leiden, The Netherlands; Laboratoire de Physique Théorique, Centre National de la Recherche Scientifique (CNRS) & Université de Toulouse - Paul Sabatier, Toulouse, France; Toulouse Institute for Infectious and Inflammatory Diseases (INFINITy), INSERM, CNRS, Université de Toulouse, Toulouse, France; Ludwig Boltzmann Institute for Rare and Undiagnosed Diseases, Vienna, Austria; Department of Dermatology, Medical University of Vienna, Vienna, Austria; Centre de Recherches sur la Cognition Animale (CRCA), Centre de Biologie Intégrative (CBI), Centre National de la Recherche Scientifique (CNRS) & Université de Toulouse - Paul Sabatier, Toulouse, France; Centre for Ecological Sciences, Indian Institute of Science, Bengaluru, India; Institute of Biology Leiden, Leiden University, Leiden, The Netherlands

## Abstract

Lymphocytes have been described to perform different motility patterns such as Brownian random walks, persistent random walks, and Lévy walks. Depending on the conditions, such as confinement or the distribution of target cells, either Brownian or Lévy walks lead to more efficient interaction with the targets. The diversity of these motility patterns may be explained by an adaptive response to the surrounding extracellular matrix (ECM). Indeed, depending on the ECM composition, lymphocytes either display a floating motion without attaching to the ECM, or sliding and stepping motion with respectively continuous or discontinuous attachment to the ECM, or pivoting behaviour with sustained attachment to the ECM. Moreover, on the long term, lymphocytes either perform a persistent random walk or a Brownian-like movement depending on the ECM composition. How the ECM affects cell motility is still incompletely understood. Here, we integrate essential mechanistic details of the lymphocyte-matrix adhesions and lymphocyte intrinsic cytoskeletal induced cell propulsion into a Cellular Potts model (CPM). We show that the combination of *de novo* cell-matrix adhesion formation, adhesion growth and shrinkage, adhesion rupture, and feedback of adhesions onto cell propulsion recapitulates multiple lymphocyte behaviours, for different lymphocyte subsets and various substrates. With an increasing attachment area and increased adhesion strength, the cells’ speed and persistence decreases. Additionally, the model can predict short-term persistent with long-term subdiffusive motility, showing a pivoting motion. For small adhesion areas, we observe that the spatial distribution of adhesions influences cell motility. Small adhesions at the front allow for more persistent motion than larger clusters at the back, despite a similar total adhesion area. In conclusion, we present an integrated framework to simulate the effects of ECM proteins on cell-matrix adhesion dynamics. The model reveals a sufficient set of principles explaining the plasticity of lymphocyte motility.

**Author summary:** During immunosurveillance, lymphocytes patrol through tissues to interact with cancer cells, other immune cells, and pathogens. The efficiency of this process depends on the kinds of trajectories taken, ranging from simple Brownian walks to Lévy walks. The composition of the extracellular matrix (ECM), a network of macromolecules, affects the formation of cell-matrix adhesions, thus strongly influencing the way lymphocytes move. Here, we present a model of lymphocyte motility driven by adhesions that grow, shrink and rupture in response to the ECM and cellular forces. Compared to other models, our model is computationally light making it suitable for generating long term cell track data, while still capturing actin dynamics and adhesion turnover. Our model suggests that cell motility is affected by the force required to break adhesions and the rate at which new adhesions form. Adhesions can promote cell protrusion by inhibiting retrograde actin flow. After introducing this effect into the model, we found that it reduces the cellular diffusivity and that it promotes stick-slip behaviour. Furthermore, location and size of adhesion clusters determined cell persistence. Overall, our model explains the plasticity of lymphocyte behaviour in response to the ECM.

## Introduction

Lymphocytes patrol in tissues and are recruited to infected areas to detect and clear the area of pathogens and cancer cells. The efficiency by which lymphocytes can find their target depends both on the characteristics of the trajectories that lymphocytes follow and the local density of the environment [1–5]. In the absence of obstacles, Lévy walks and persistent random walks outperform Brownian walks. Lévy walks are characterized by long strides in their trajectories such that they cover larger areas than Brownian walks. In environments crowded with obstacles Brownian walks perform better, as the more compact trajectory leads to more thorough local exploration [1]. Even more local exploration follows from subdiffusive random walkers, which diffuse less far than could be expected from their speed and persistence. Consistent with these characteristics, in the densely populated lymph nodes T lymphocytes perform Brownian walks [6, 7] or persistent random walks [8]. In brain tissue, T cells perform Lévy walks [4]. In pancreatic islets, CD4+ T cells perform subdiffusive random walks, whereas CD8+ T cells perform confined random walks [9]. The characteristics of these different types of motion, including speed distribution and mean squared displacement (MSD), determine how efficiently lymphocytes can find their targets *in vivo*. Therefore, it is key to understand what factors give rise to these different types of walks.

The plasticity of lymphocyte motility behaviour is dictated both by environmental factors and by cell intrinsic features [10, 11]. An *in vitro* study has shown that the type of extracellular matrix (ECM) used as cell culture substrate affects the motility patterns of B lymphocytes, possibly due to the attachment strength [12]. On fibronectin, B lymphocytes show higher diffusivity and more effective displacement than on collagen IV substrates where cells move more slowly. The B lymphocytes form larger adhesive connections with fibronectin than with collagen IV, and on fibronectin the cells change shape more rapidly than on collagen IV. Similar effects have been found for T lymphocytes. The majority of cells on a casein substrate move through multiple, distinct and temporary adhesion zones, i.e., walking motility, whereas on ICAM-1 substrates, the majority of cells make one continuous contact zone with the substrate, i.e., sliding motility [13]. Apart from these environmental effects, cells also show large individual variation in their motility patterns. On fibronectin, individual B lymphocytes showed either floating-like behaviour with little attachment, dynamic attachment leading to stepping/walking behaviour, or sustained attachment leading to cells pivoting around their adhesive area [12]. Similarly, T lymphocytes showed either walking, mixed or sliding behaviour, with frequencies depending the type of culture substrate [13].

It is still poorly understood what causes, on the one hand, the consistent differences in motility modes between culture substrates, and on the other hand, the large individual differences between cells on the same type of substrate. To answer this question, we propose a simplified mathematical model of cell motility and the adhesive interaction with the ECM. Modelling can help in understanding which elements in cell-matrix interactions are important for the variety in cell motility [14], and previous theoretical studies have already provided useful insights into this problem. Copos et al. [15] asked what causes the cellular extension and retraction cycles driving the motility of *Dictyostelium discoideum* cells. They modelled *D. discoideum* movement in a force-based model. Depending on the density of adhesion binding sites in the substrate, or the strength of these adhesions, the model displayed different motility types. For low densities of adhesion binding sites in the substrate and low adhesion strength, gliding motility was observed. Increasing the density of the binding sites or the adhesion strength led to a stepping motility mode of reduced speed. For the highest adhesion densities or adhesion strengths, the cells became stationary. Thus, the cells moved faster in the gliding motility mode than in the stepping motility mode. Although this agreed with preliminary experimental results on *D. discoideum* cells [15], these results contradict observation in lymphocytes: T cells move faster in stepping motility mode than in gliding motility mode [13]. Furthermore, as a one-dimensional model, it cannot produce two-dimensional cell tracks and it is computationally too heavy for producing the large amounts of cell track data required for our purpose. Phase-field models have been used for studying the influence of the ECM on cell motility in two dimensions [16, 17]. Ziebert et al. [16] present a phase-field model to study how actin polymerization, dynamical adhesion site formation, and substrate compliance together determine the characteristics of cell trajectories. Simulated cells displayed a gliding motion when substrate stiffness was high, the protrusion strength was large and adhesions formed at a high rate. At intermediate substrate stiffnesses with sufficiently high protrusion strength and intermediate adhesion formation, the cells displayed a stick-and-slip motion. Thus, phase-field models have provided key insights into how the extracellular matrix affects cell motility. However, their high computational costs currently limit the production of the large data volumes required for statistical analyses of cell trajectories. A first step in this direction was taken by Yu et al. [18] who have introduced a computationally efficient, coarse-grained model to study long term cell persistence. The model considered spheroid cells with a fixed pool of focal adhesions. These adhesions were assumed to be widely dispersed within the cells for soft substrates and more narrowly dispersed for rigid substrates. The increased persistence times on rigid substrates led to durotaxis, i.e., preferential movement towards stiffer substrates. In their model, Yu et al. [18] assumed a direct dependence of cell persistence on adhesion distribution. In our work, we hope to study how the adhesion distribution emerges *de novo* from the interaction between adhesion formation, adhesion detachment and cell motility.

Thus, previous models are either too computationally expensive or do not model the effect of adhesion on the microscopic level. The Cellular Potts model is conceptually closely related to phase-field models, but is computationally much lighter. The Act model [19], a recent extension of the Cellular Potts model, provides of phenomenological model of actin dynamics. Interestingly, this model can already display multiple motility modes: Simulated cells show intermittent (stop-and-go) or persistent random walks. An in-depth characterization revealed that the model displays universal coupling between speed and persistence, and specifically that speed increases linearly with protrusion strength, whereas persistence time increase exponentially with protrusion strength [20].

Here, we extend the Act model with cell-ECM interactions. The model combines coarse-grained actin dynamics, with simplified dynamics of adhesion turn-over and detachment, resulting in a diverse palette of cell motility. In a second version of the model, the cell-ECM adhesions promote cell protrusion by inhibiting retrograde actin flow. Our model can simulate cell motion with sufficient detail on the location and size of adhesive patches, while being computationally light enough for statistical analysis of cell motility. With the actin component and cell-matrix adhesion component of the model, we are able to reproduce a variety of cell motion types, similar to the behaviour seen in other models that also include those two components [15–17]. In addition to persistent random walks and ballistic cell motility, the extended model can also predict anomalous diffusion with long-term subdiffusive behaviour, showing all three phases of lymphocyte motility on fibronectin found in the experimental work by Rey-Barroso et al. [12]. Our model shows that simple cell-ECM interactions can drastically alter cell motility. Thus, adhesion dynamics can play a key role in the plasticity of motility in response to ECM composition.

## Results

In this section, we first introduce our extension of the Act-model with cell-matrix adhesions. We then show that this model can reproduce a wide range of lymphocyte motility modes. Next, we extend the model with feedback from the adhesions onto the actin polymerization force and show that we can capture more dynamic motility behaviours. Overall, our model recapitulates the diversity of lymphocyte motility modes and provides insight into the mechanisms underlying such behavioural diversity.

### Modelling cell-matrix adhesions

Our computational model is based on the Act model [19, 20], an extension of the Cellular Potts model (CPM, [21, 22]) for efficient modelling of persistent, amoeboid cell motility emerging from feedback mechanism inspired by detailed insights into actin-polymerization driven cell motility. Depending on its parameter settings, the model produces a variety of realistic cell shapes, cell polarization, and cell trajectories. In short, this extension keeps track of recent “actin polymerization” through *Act values* at each lattice site. Novel protrusions get a high *Act value* and cell protrusions at sites with locally high *Act values* are favoured. Two important parameters for this are *λ*_*Act*_, the weight of the Act model that can be interpreted as the maximum protrusive force of the actin network, and Max_*Act*_, the maximum Act value, interpretable as the lifetime of an actin subunit within the actin network (see Table 1). In addition, our model takes cell-matrix adhesion into account and reflects the dynamic processes of such adhesions (Fig 1). We will shortly explain the addition of cell-matrix adhesions below. For further details, see the Methods section.

**Table 1.** List of parameters involved in adhesion dynamics and values used for simulations.

| Parameter | Description | Values |  |  |
| --- | --- | --- | --- | --- |
|  |  | Figs 2,3,4 | Figs 6,7 | Fig 8 |
| $\lambda_{Act}$ | Weight of the Act-extension, the maximum protrusive force induced by actin polymerization | 240 | 120, 240 | 240 |
| $\text{Max}_{Act}$ | Maximum value of the Act-field, actin lifetime | 120 | 120 | 120 |
| - | Act-value threshold above which adhesion formation is possible | 0.75 | 0.75 | 0.75 |
| $p_s$ | New adhesion formation rate | 0.004-0.020 | 0.001-0.004 | 0.003,0.001 |
| $p_e$ | Rate of neighbouring grid site to become adhesion site if not already so. | 0.0055 | 0.0055 | 0.0015,0.004 |
| $p_d$ | Rate of unbinding adhesion site dependent on adhesionless neighbouring sites | 0.0008 | 0.001 | 0.0004 |
| $\lambda_{adh}$ | Energy required to rupture adhesion upon retraction of the cell | 20-100 | 20-100 | 60 |
| $f$ | Prefactor for the adhesion feedback onto Act model | - | $b-1$ | $b-1$ |
| $b$ | Base value of $f$ in absence of adhesions | - | 0.5 | 0.5 |
| $s$ | Adhesion area fraction saturation threshold above which $f = 1$ | - | 0.1 | 0.12 |

**Fig 1.**
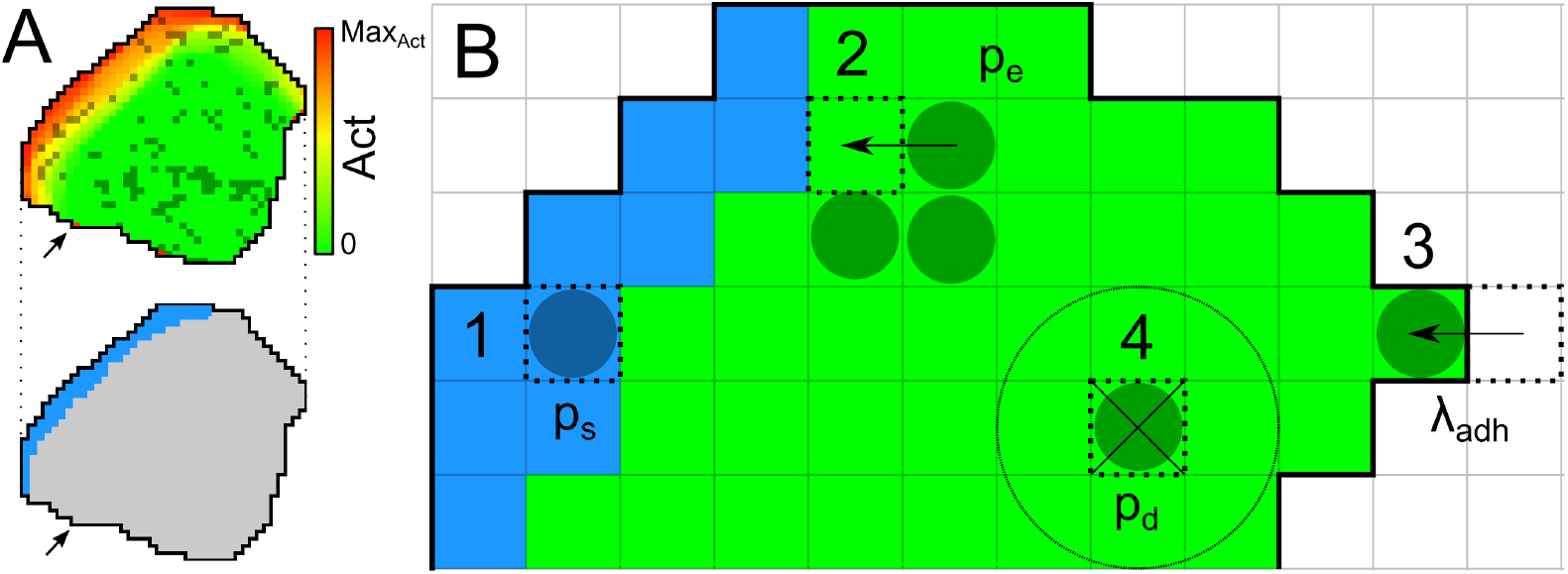
Overview of the adhesion processes within the model. A) Top: overview of a simulated cell. Red to yellow shading indicates the Act-level of each lattice site according to the colourbar. Darker coloured lattice sites contain an adhesion. Bottom: the same cell with in blue the region where new adhesions can form, as the local Act-levels exceeds the 0.75 Max_*Act*_ threshold. Both: Arrows point to area with one lattice site with high Act-level due to a recent extension of the cell (top, red), but the geometric mean of Act-levels does not exceed the threshold and hence new adhesions cannot form there (bottom, grey). B) Visual summary of adhesion processes. Dark coloured circles indicate lattice sites containing an adhesion. 1) New adhesions can form spontaneously with probability *p*_*s*_ at cell lattice sites where the geometric means of Act values exceeds the threshold of 0.75 Max_*Act*_ (blue region). 2) An adhesion patch can grow by Eden growth. A random neighbour of an adhesion site is selected. When it does not contain an adhesion yet, the patch extends into that lattice site with probability *p*_*e*_. 3) To break an adhesion, retractions must overcome an energy cost Δ*H* = *λ*_*adh*_. 4) Adhesions can also unbind spontaneously, depending on the number of neighbouring lattice sites without adhesions and probability *p*_*d*_.

Cell-matrix adhesions are represented at a subcellular level as discrete entities: A subcellular CPM lattice site can either contain an adhesion or not. Lattice sites are approximately 600 nm wide, considering that simulated cells contain approximately 1000 CPM lattice sites each, and that B cells and T cells on a substrate cover an area of approximately 360 µm^2^ [12, 23]. Observations show that single adhesion units in lymphocytes are approximately 100 nm in diameter [23], so a single adhesion lattice site in the model represents a small number (≤5) of adhesion units, considering adhesion unit density of 5 clusters/µm^2^ [23].

The formation of new adhesions depends on actin polymerization, membrane protrusion and the distribution of integrins at the leading edge of the cell [24–26]. As actin activity and the leading edge are marked in the Act model by lattice sites with high Act values, we let new adhesions appear at lattice sites with a locally high Act-level. More precisely, defining 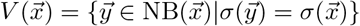 as the Moore neighbourhood of lattice site 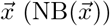 restricted to the same cell as site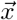, a lattice site 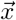 receives an adhesion with probability *p*_*s*_ (Fig 1A and 1B, process 1) if the geometric mean of Act-value 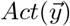, with 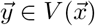 exceeds a threshold 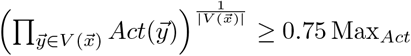 ≥0.75 Max_*Act*_. Here, *p*_*s*_ lumps together the effect of multiple biological processes, including the rate at which integrin molecules bind to their ligands in the extracellular matrix, and the concentration of integrins at the cell front.

Once an adhesion has formed, it can either expand into an adhesion patch, or unbind. Biologically, adhesion patches expand radially with some bias in the direction of the cell front [13]. A detailed biophysical model suggests that local integrin-substrate bonds favour the formation of adjacent bonds, because they stabilize the membrane thus reducing membrane fluctuations [27]. We simplify patch expansion phenomenologically using the Eden model [28] of radial colony growth. Empty lattice sites adjacent to an existing adhesion site join the adhesion patch with probability *p*_*e*_ (Fig 1B, process 2).

Cell-matrix adhesions unbind due to cellular forces or spontaneously. Adhesion patches rupture from the edges of the patches [13]. We model this rupture only at the edge of the cell, where contraction forces of the cell can break bonds. Integrin bonds are known to show catch-slip bond behaviour, meaning that initially the bond strengthens with increase of force, but will still break if enough force is applied. Here, we neglect this specific behaviour and associate a single required energy cost of *λ*_*adh*_ with the rupture of adhesions at the retracting edge (Fig 1B, process 3). *λ*_*adh*_ is given by the binding affinity between integrins and their ligands and the concentration of integrins bound to the ECM. In addition, we assume that association with adjacent adhesions reduces the spontaneous unbinding rate. The probability that an adhesion site unbinds thus becomes 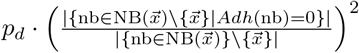, where 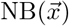 indicates the Moore neighbourhood of lattice site 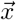 (Fig 1B, process 4).

All in all, the proposed model extension for adhesions is relatively simple and computationally light. All adhesion dynamics are governed by the four parameters *p*_*s*_, *p*_*e*_, *p*_*d*_ and *λ*_*adh*_. An overview of all the relevant parameters is shown in Table 1.

Estimates of length scale, time scale, and the parameter Max_*Act*_, as well as the relative magnitudes of *λ*_*Act*_ and *λ*_*adh*_ can be found in S2 Material.

### Model reproduces crawling and pivoting motility

In this section we first investigate the influence of the adhesion formation probability *p*_*s*_, and the adhesion strength *λ*_*adh*_. Figure 2(A-D) summarize the variety of patterns observed in the model. If *λ*_*adh*_ is low to moderate (Fig 2A,2C, S1 Video) cells show crawling motility as in the standard Act-model [19]. For larger values of *λ*_*adh*_ cell show less persistent motility (Fig 2B,2D, S1 Video). For high values of *λ*_*adh*_ and *p*_*s*_ cells show pivoting motility. For these parameter values, the adhesions form easily and require much energy to break, such that cells will remain stuck in place on the matrix. However, they can still make protrusions around them, leading to pivoting motility (Fig 2D, S1 Video).

**Fig 2.**
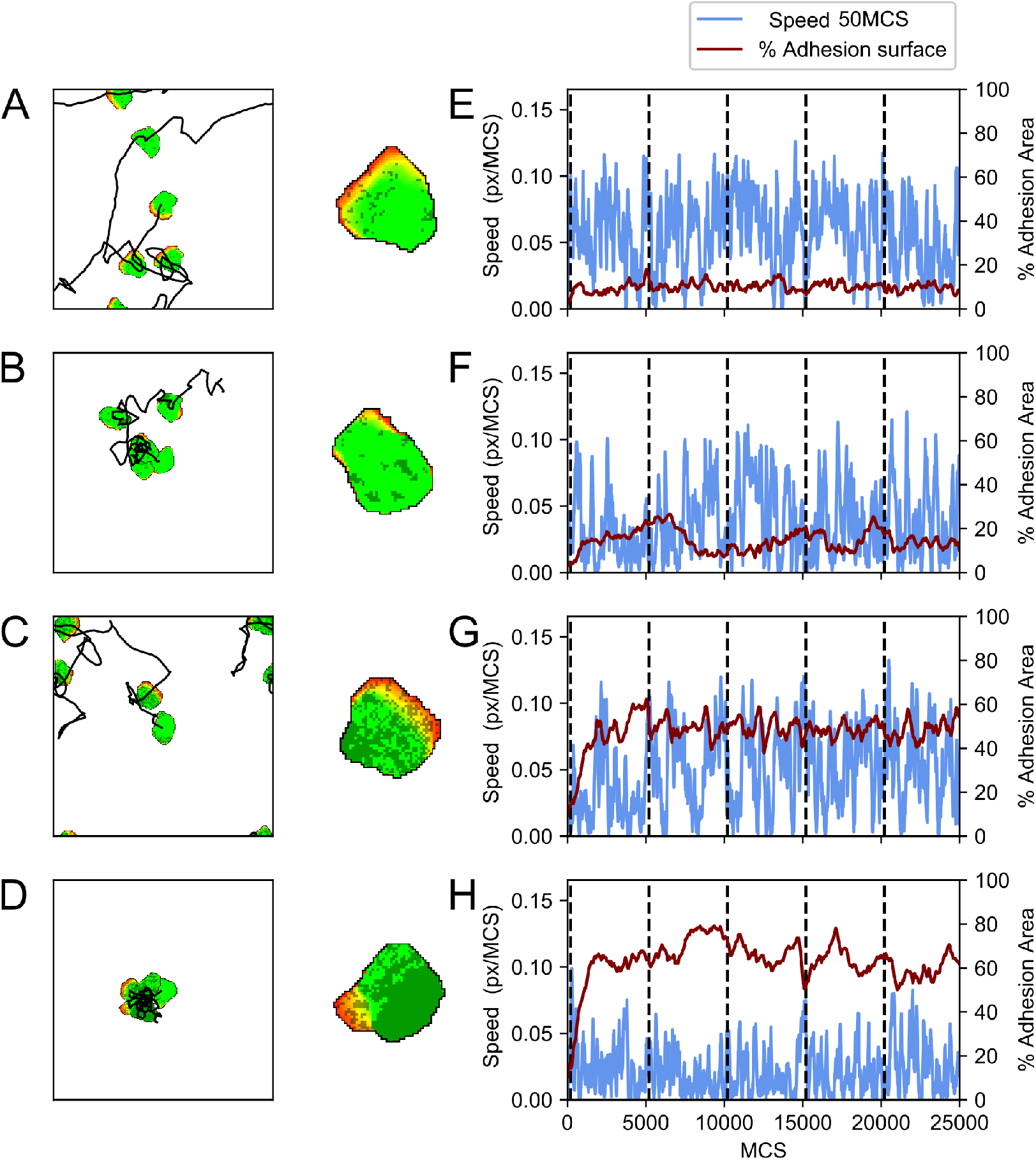
Simulations of the model showing different motility types. A-D) First column, a display of a single cell at 5000 MCS interval snapshots combined with the cell centre’s trajectory. Each trajectory starts in the centre of the field and periodic boundaries are used. Second column, a close-up of the cell with the adhesions displayed in a darker colour. E-H) A plot of the cell’s speed and percentage of the cell’s area containing adhesions corresponding to the track on the left. Vertical dashed lines indicate the times of the snapshots on the left. Parameters are: A,E) *λ*_*adh*_ = 20, *p*_*s*_ = 0.004, B,F) *λ*_*adh*_ = 100, *p*_*s*_ = 0.004, C,G) *λ*_*adh*_ = 20, *p*_*s*_ = 0.02, D,H) *λ*_*adh*_ = 100, *p*_*s*_ = 0.02. Furthermore, *p*_*d*_ = 0.0008 for all cases. These simulations are also available as S1 Video.

To further characterize the cell motility and the effect of the adhesion patches, we measured the cell speed alongside with the summed area of the adhesions (Fig 2(E-H). Comparing these four parameter settings (Fig 2), *p*_*s*_ regulates the adhesion area. At relatively low values of *p*_*s*_ = 0.004 (Fig 2E,F), the cells form low adhesion area, and at higher values of *p*_*s*_ = 0.02, we observe a larger adhesion area (Fig 2(G,H). *λ*_*adh*_ regulates the cell speed: at relatively low values of *λ*_*adh*_ = 20 (2E,G) the speed fluctuates less than at higher values of *λ*_*adh*_ = 100 (2F,H). Interestingly, in (Fig 2B we observe stick-slip-like behaviour: upon a reduction of the adhesion area, the speed increases. Overall, the behaviour of our model resembles observations by Rey-Barroso et al. [12] that B cells on fibronectin with fluctuating adhesion areas showed walking behaviour, and cells with large and sustained adhesion surfaces displaced very little and the adhesion patch did not displace. The behaviour of the model also agrees with observations of Jacobelli et al. [13] that T cells displaying a gliding motion with higher adhesion area have lower speed than cells with a walking motion with lower adhesion area.

### Adhesions slow down cell motion and reduce diffusivity

The examples shown in Fig 2 and S1 Video suggest that higher adhesion area correlates with reduced cell speed and reduced cell diffusivity. To characterize this potential correlation, we analyse 1000 independent runs for different combinations of *p*_*s*_ and *λ*_*adh*_ measure the average cell speed and adhesion area. Fig 3A shows the mean speed of the cells as a function of *p*_*s*_ and *λ*_*adh*_, and Fig 3B shows the diffusivity. In both panels, symbols with the same value of *λ*_*adh*_ are connected and symbols of the same colour have the same value of *p*_*s*_. Fig 3A shows that increasing the value of *λ*_*adh*_ decreases cell speed and that the average adhesion area increases with the value of *p*_*s*_. Furthermore, for high values of *λ*_*adh*_, the effects of *p*_*s*_ on cell adhesion are larger, and for high values of *p*_*s*_, the effects of *λ*_*adh*_ on cell speed are also larger. Fig 3B plots the diffusivity of the cells and adhesion area as a function of *p*_*s*_ and *λ*_*adh*_. Diffusivity was obtained by measuring the slope of mean squared displacement curves from 11500 to 24000 MCS. Similar to the cell speed, increasing *λ*_*adh*_ results in lower diffusivity and the effect is larger for higher values of *p*_*s*_. Moreover, the effect of *λ*_*adh*_ is larger on cell diffusivity than on cell speed. The drop in instantaneous cell speed (from highest to lowest in Fig 3A about 50% smaller) is modest compared to the drop in diffusion coefficient (Fig 3B, about 380% smaller). Cell diffusivity is determined by instantaneous cell speed and persistence of movement direction. Thus, this observation suggests that the drop in diffusivity is for a large part caused by a loss of cell persistence.

**Fig 3.**
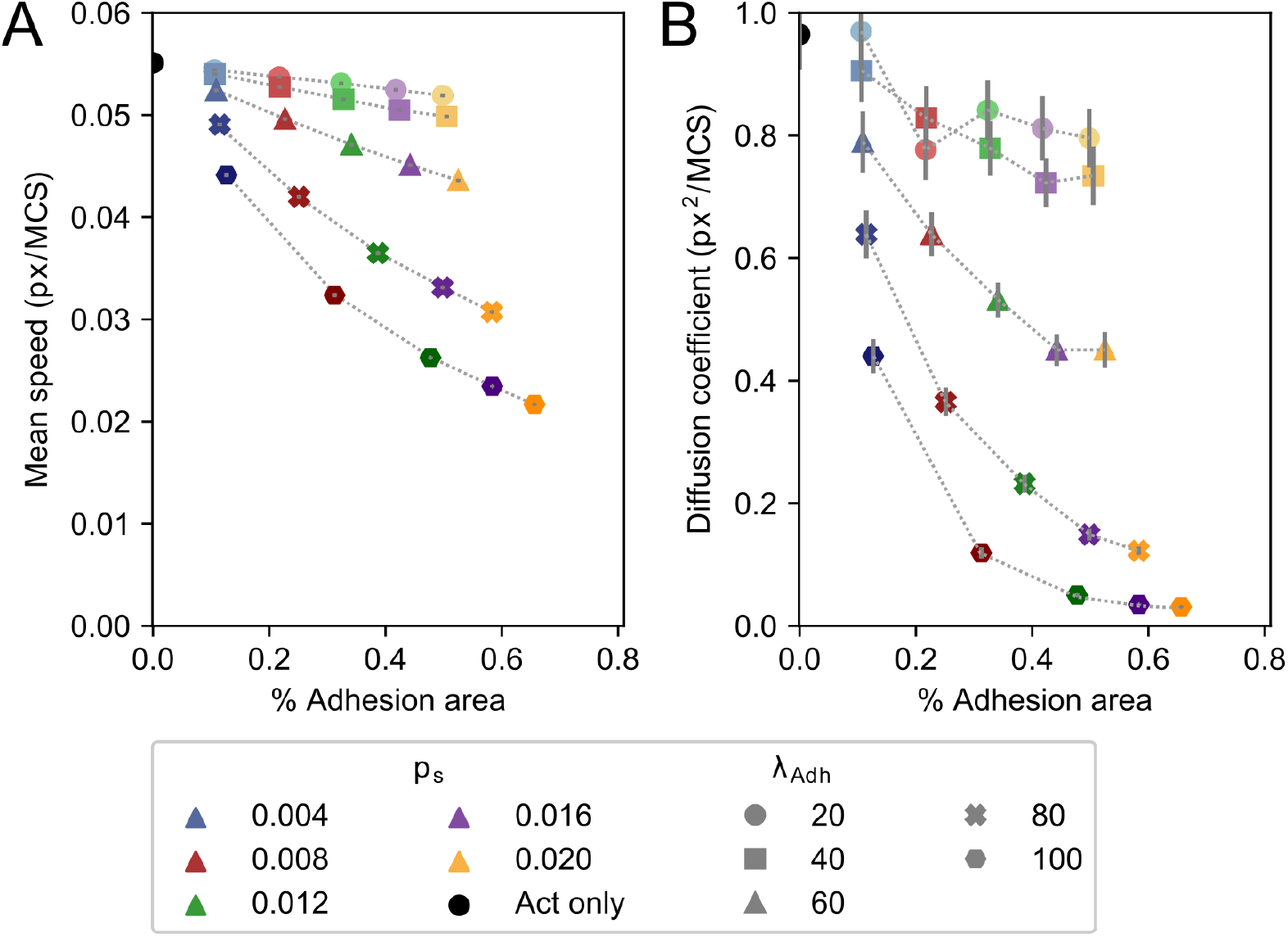
Mean speed, diffusion coefficient, and mean adhesive area change by increasing *p*_*s*_ and *λ*_*adh*_. Mean speed (A) and diffusion coefficient (B) plotted against mean percentual adhesion area for different values of parameters *p*_*s*_ and *λ*_*adh*_. Each dot represents the mean of 1000 independent simulations. Different colours indicate different *p*_*s*_, where shades from light to dark and marker symbol indicate *λ*_*adh*_ ∈ {20, 40, 60, 80, 100}. For reference, the mean speed and diffusion coefficient of the Act model without any adhesions are indicated by a black circle. Error bars indicate 95% confidence intervals.

### Adhesive cells’ reduced diffusivity is due to reduced persistence times

To further characterize the cause of the reduced diffusivity at high values of *p*_*s*_ and *λ*_*adh*_, we fitted the mean squared displacement curves with a persistent random walker model, the Fürth equation [29, 30]:

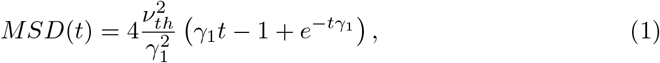

where *ν*_*th*_ is the walker’s speed and *γ*_1_ is its persistence time (S1 Figure). The persistent random walker model fits well with the MSD curves at long time scale, but it fails at the short time scale where the cell trajectories are mainly determined by the random fluctuations in the CPM. We therefore extended the Eq 1 with a term for translational diffusion [31],

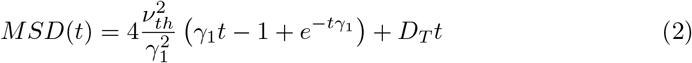

The extended Fürth equation (Eq 2, S1 Figure) gives good fits for most cases (Fig 4A-C). The persistence times, as obtained from the extended Fürth equation (Eq 2), indicate that indeed the reduced diffusivity can be attributed for a large part to a reduced persistence time with higher adhesion energies and large adhesive areas (S2 Figure). However, for the highest values of *λ*_*adh*_ = 80 to *λ*_*adh*_ = 100 Eq 2 still does not fit well with the data (Fig 4D). We attempted to improve the fit by increasing the initialization period left out of the MSD computation, in order to compute the MSD of cells closer to their dynamic equilibrium in both Act model dynamics as well as adhesion-extension dynamics. This barely improves the fit and suggests that cell motion in this regime cannot be correctly described by a persistent random walker with translational diffusion.

**Fig 4.**
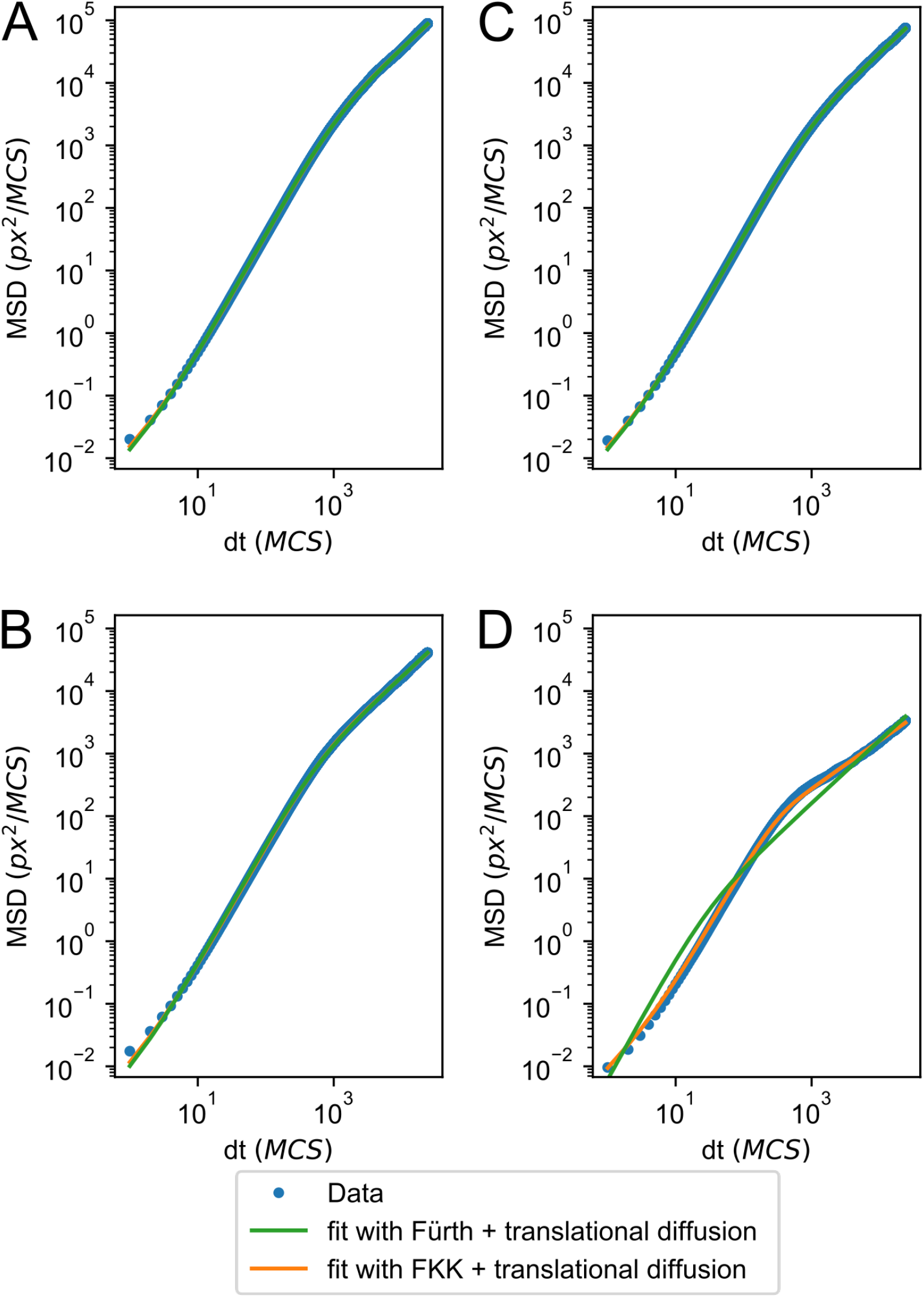
MSD fits either a persistent random walker or a subdiffusive persistent random walker. Log-log plot of MSD for the four scenarios in Fig 2, with fits of Eqs 2 and 4. Parameters are: A) *λ*_*adh*_ = 20, *p*_*s*_ = 0.004, B) *λ*_*adh*_ = 100, *p*_*s*_ = 0.004, C) *λ*_*adh*_ = 20, *p*_*s*_ = 0.02, D) *λ*_*adh*_ = 100, *p*_*s*_ = 0.02.

### Strongly adhesive cells show subdiffusive behaviour

Previously the fractional Klein-Kramers (FKK) process was proposed to describe the motility of transformed Madin-Darby canine kidney cells [32],

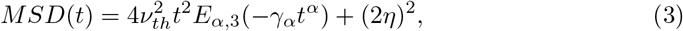

where *E*_*α*,3_ is the generalized Mittag-Leffler function and *η* is a noise term. The standard Fürth model (Eq 1) is a special case of the FKK-process for *α* = 1 and *η* = 0, where *α* parameterizes the long-term diffusive behaviour. Since we already concluded that translational diffusion plays a significant role in the short-time scale of the CPM, we assume that the noise term is due to translational diffusion, thus obtaining,

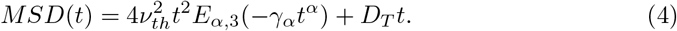

which reduces to Eq 2 for *α* = 1. For *t* →∞, Eq 3 and Eq 4 can be approximated by 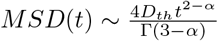 [32]. So for *α >* 1, the long-term behaviour is subdiffusive, whereas for *α <* 1, the long-term behaviour is superdiffusive.

In the cases where Eq 2 fits well, we obtain *α ≈*1 (Table 2). However, for the cases were Eq 2 fits badly, Eq 4 has a better fit and *α >* 1 (Fig 4, Table 2) indicating subdiffusive behaviour. This behaviour corresponds to cases where the cells are strongly attached to the matrix, pivoting around their adhesion patch. Thus, these cells move persistently on a local scale as they have a single protrusion front, but they move subdiffusively on a longer timescale as the stay within a confined area.

**Table 2.** Fitted values of *α* from Eq 4 for different values of *λ*_*adh*_ and *p*_*s*_.

| Parameters | $\alpha$ |
| --- | --- |
| $\lambda_{adh} = 20, p_s = 0.004$ | 1.019 |
| $\lambda_{adh} = 20, p_s = 0.020$ | 1.013 |
| $\lambda_{adh} = 100, p_s = 0.004$ | 1.024 |
| $\lambda_{adh} = 100, p_s = 0.020$ | 1.257 |

### Feedback of adhesions onto propulsion efficiency

So far, the model can explain (i) sliding and stepping, and (ii) pivoting cell behaviour as observed in Ref. [12]. However, the current model still fails to explain (iii) the floating phase that was also observed in Ref. [12], i.e., the observation that B cells with a low adhesive area or no adhesive area on a fibronectin substrate show low displacement compared to cells with dynamic attachment [12].

To explain also such floating behaviour, we extended the model as follows. In presence of a stable adhesion, the forces generated by actin polymerization are transferred onto the matrix leading to protrusion. With reduced adhesions, actin polymerisation more often leads to treadmilling. Thus, in presence of cell-matrix adhesion, actin polymerisation more efficiently translates to cell protrusion [33–35]. We model this effect using a prefactor *f* to *λ*_*Act*_, such that it dynamically alters the strength by which the Act-values affect the cell protrusions, and hence the protrusion force. For simplicity, the protrusion efficiency increases linearly with the cell’s total adhesion area. Furthermore, we assume that there is a threshold adhesion area at which the adhesive force suffices to withstand the forces of actin polymerization. Hence, we define:

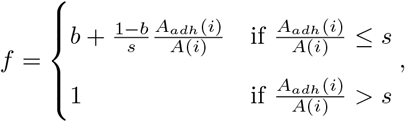

with *b* the baseline protrusion efficiency and *s* the threshold adhesion area. Thus the effect of the adhesion area on the propulsion strength only affects cell motility if the adhesion area is below or near the threshold *s*. A schematic overview is shown in Fig 5.

**Fig 5.**
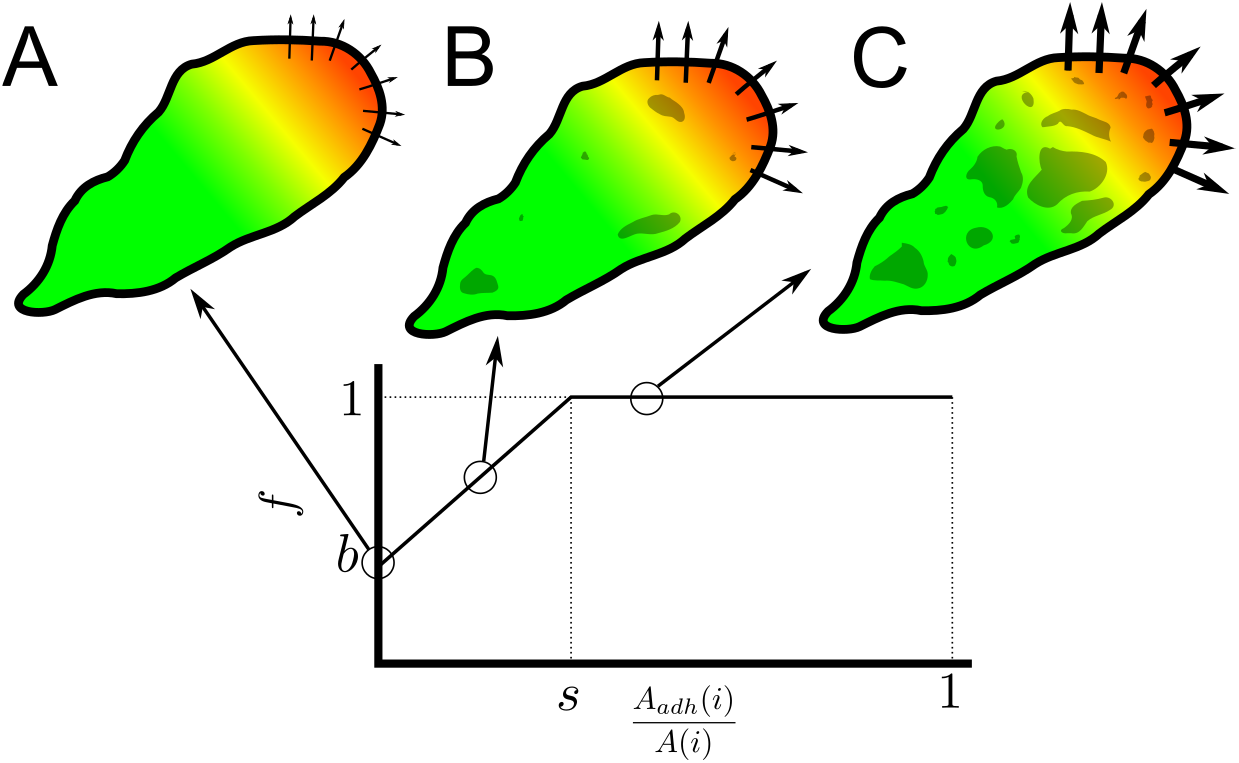
Schematic representation of the effect of adhesion on cell propulsion strength. Colour schemes are similar to Fig 1A. Arrow width corresponds to the effective protrusion strength *fλ*_*Act*_. A) In the absence of adhesions, the propulsion prefactor *f* is equal to the base level *b*. B) Below the saturation point *s, f* increases linearly with adhesion area. C) Above the adhesion area saturation point *s*, prefactor *f*, and thus effective protrusion strength *fλ*_*Act*_, are maximal.

### Extended model reproduces all three phases of cell motility

To test if the new model indeed reproduces floating motility alongside the other two phases of motility observed in Ref. [12], we focus on parameter combinations that result in adhesion areas below or around the threshold *s*. For adhesion areas above the threshold *s*, the model behaviour does not change. We choose *s* = 0.1 and a baseline protrusion efficiency *b* = 0.5. From the previous section, we know that *p*_*s*_ is the main parameter controlling adhesion area, so we chose *p*_*s*_ ≤0.004.

The model shows a variety of behaviours depending on the value of *p*_*s*_ (Fig 6, S2 Video). For very low values of *p*_*s*_ = 0.001 (Fig 6A), cells make only a small number of tiny adhesion patches and thus have a small adhesive area. Furthermore, they explore relatively small areas (Fig 6A). Nevertheless, their trajectories can still be described well with Eq 2 or with Eq 4 with *α* = 0.974, so the type of motion can still be classified as a persistent random walk, albeit with lower persistence time (S3 Figure). For *p*_*s*_ = 0.004 (Fig 6B), the mean adhesive area approaches the threshold. In this case the diffusivity and persistence are lower compared to the model without the adhesion-protrusion feedback, but the cell speed is comparable (Figs 6E,7A). In between these two adhesive regimes, stick-slip behaviour is observed (Fig 6C). Interestingly, in contrast to the initial model, the cells accelerate as the adhesion areas grow and they decelerate when they have lost their adhesions (Fig 6F).

**Fig 6.**
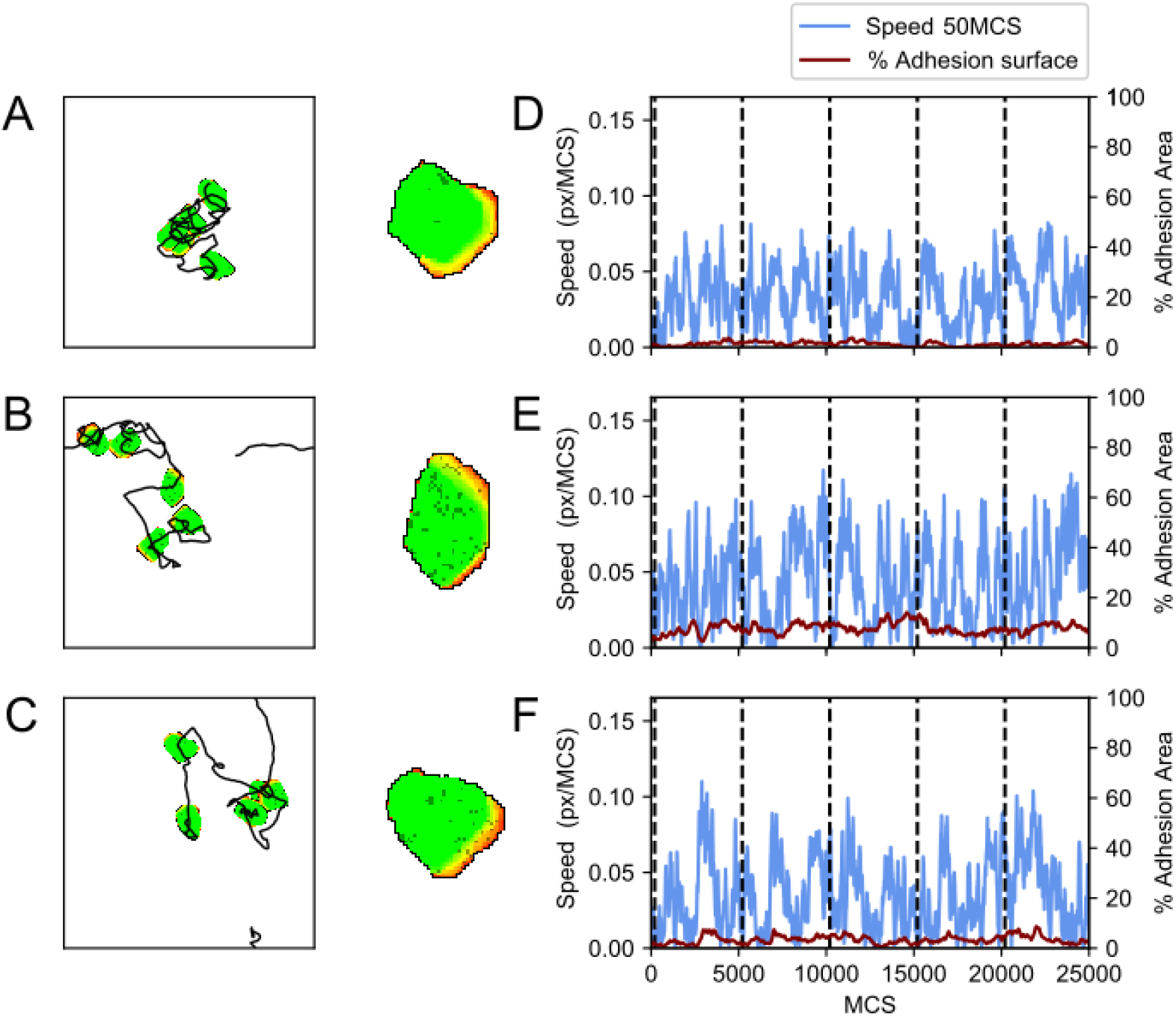
Simulations of the model with adhesion-propulsion feedback. A-C) First column, a display of a single cell at 5000 MCS intervals combined with trajectory of the cell centre. Each trajectory starts in the middle and periodic boundaries are used. Second column, a close-up of the cell with the adhesion displayed in a darker colour. D-F) A plot of the cell’s speed and percentage of the cell’s area containing adhesions corresponding to the track on the left. Parameters are: A,D) *λ*_*adh*_ = 100, *p*_*s*_ = 0.001, B,E) *λ*_*adh*_ = 100, *p*_*s*_ = 0.004, C,F) *λ*_*adh*_ = 100, *p*_*s*_ = 0.0025. Furthermore *p*_*d*_ = 0.008 for all cases. These simulations are also available as S2 Video.

Figure 7 shows the average speed (Fig 7A) and diffusivity of cells (Fig 7B) with low adhesive areas for both the initial and the extended model. The cell speed and cell diffusivity of the standard Act model are shown in the figure for reference. The cell speed and cell diffusivity of the initial model follow the same trend as observed in Figure 3, both decreasing for increasing values of *p*_*s*_ and *λ*_*adh*_. The model converges to the Act model for low adhesion areas as expected (*λ*_*Act*_ = 240 (black cross)). Similarly, the extended model converges to the Act model for low adhesion area (black plus sign), where *λ*_*Act*_ = 120 corresponds to the baseline propulsion strength *b* = 1*/*2. Near the adhesion area threshold of *s* = 0.1 both models show the same behaviour. The mean speed and diffusion coefficient increase monotonically between these two extremes: higher adhesion areas lead to higher speed. Whereas the value of *λ*_*adh*_ has little effect on the cell speed in this parameter regime, *p*_*s*_ determines the adhesion area and thus has a large effect on cell speed. Remarkably, there is a slight difference in mean adhesion area between the models. This small effect is likely due to the positive feedback loop between adhesion growth and cell propulsion in the extended model.

In conclusion, by adjusting the parameters *p*_*s*_ and *λ*_*adh*_, the extended model provides a minimal model explaining the three motility phases of B cells observed on fibronectin: for low cell-matrix attachment the cells have reduced motility (floating motility), for increased cell-matrix attachment the cells form dynamic adhesion areas and display increased motility (sliding-stepping motility), and for the strongest cell-matrix attachment the cells displayed sustained attachment with low displacement (pivoting motility) [12].

**Fig 7.**
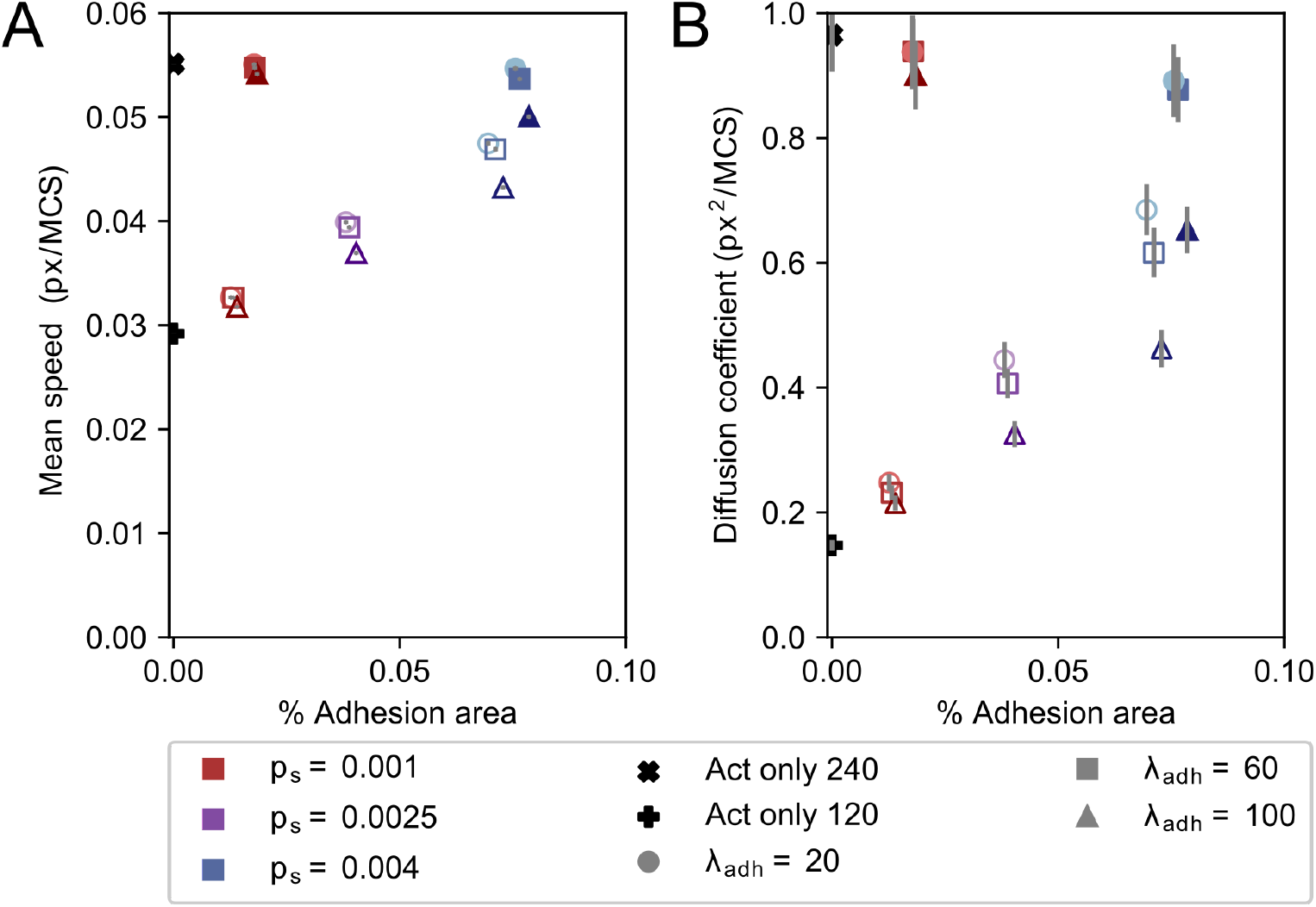
Mean speed, diffusion coefficient and adhesion area differ between the model with and without adhesion-propulsion feedback. Mean speed (A) and diffusion coefficient (B) plotted against mean percentual adhesion area for different values of parameters *p*_*s*_ and *λ*_*adh*_. Each dot represents the mean over 1000 independent simulations. Filled symbols are results from the model without adhesion-propulsion feedback, and empty symbols show results from the model with adhesion-propulsion feedback. Different colours indicate different *p*_*s*_, where shades from light to dark and marker symbol indicate *λ*_*adh*_ ∈ {20, 60, 100}. Error bars indicate 95% confidence intervals.

### Adhesion growth dynamics change persistence time

So far, we have mostly looked the effect of adhesions formed at the cell front. However, the distribution of adhesive patches over the cell may have a large effect on cell motility patterns. For example, the distribution of the adhesion clusters differs between walking and sliding T cells [13]. Sliding T cells have a large contact area at the cell front. Walking T cells, in contrast, had multiple distinct adhesion areas that were distributed over the cells, including the rear end of the cell.

To gain further insight in the impact of adhesion patch distribution on cell motility, we also explored parameter settings resulting in the same adhesion area but with different adhesion cluster size distributions, by varying the formation rate for new adhesions (*p*_*s*_) and adhesion growth rate (*p*_*e*_). Fig 8 shows the results of two such parameter settings resulting in the same average adhesion area. The blue parameter set obtains its adhesive area mostly through the formation of new adhesions (*p*_*s*_ *> p*_*e*_, Fig 8A: blue), whereas the other more rapidly grows out adhesion clusters (*p*_*s*_ *< p*_*e*_, Fig 8A: orange, see also S3 Video). This results in different cluster size distribution (Fig 8B), with only small clusters when *p*_*s*_ *> p*_*e*_ (blue line), and small clusters combined with a few large ones when *p*_*s*_ *< p*_*e*_ (orange line).

**Fig 8.**
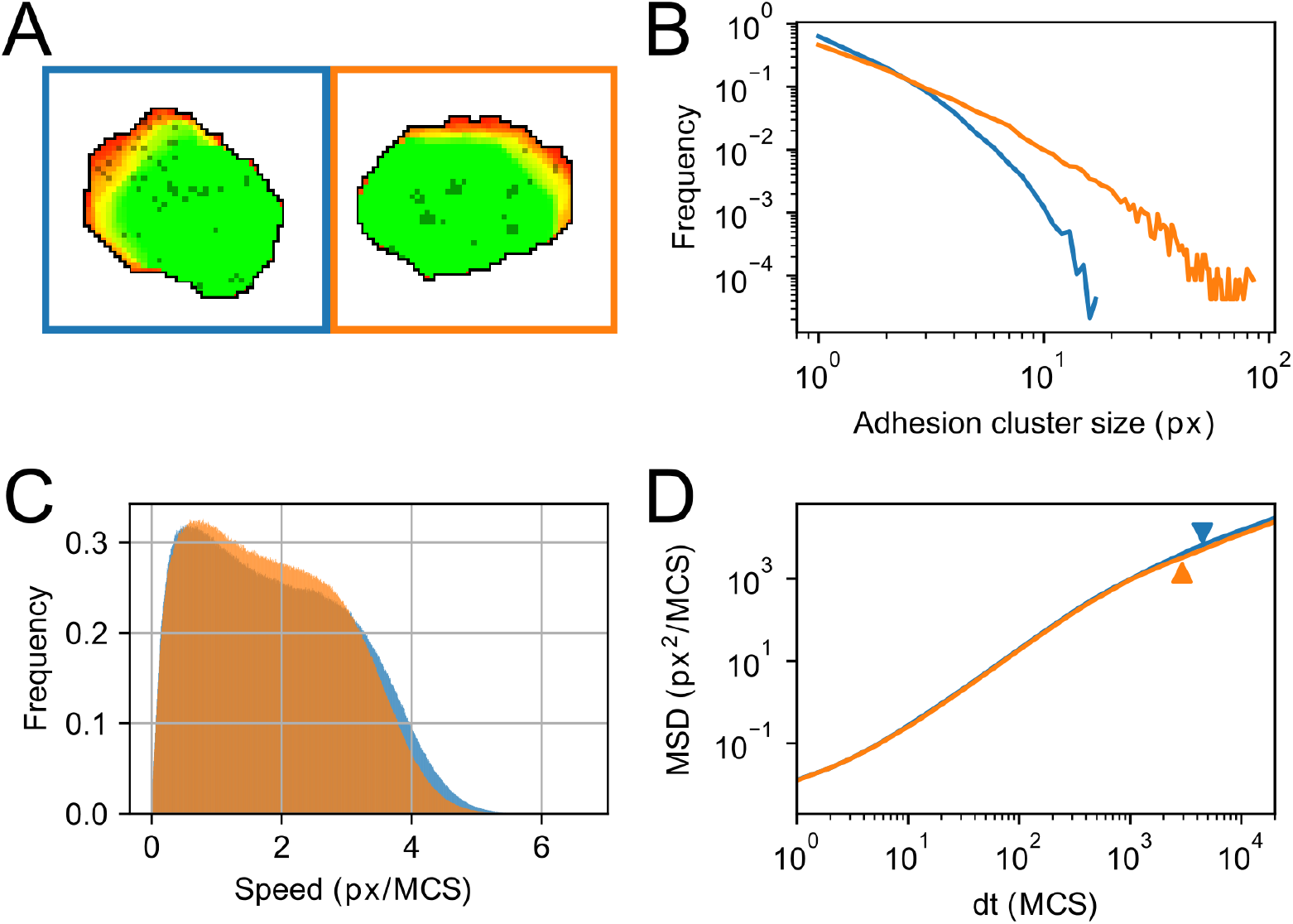
Adhesion growth dynamics influence adhesion cluster size, cell speed and MSD. (A) Example of different adhesion cluster size for different parameter values of *p*_*s*_ and *p*_*e*_. Blue: *p*_*s*_ = 0.003, *p*_*e*_ = 0.0015 resulting in multiple small adhesions. Orange: *p*_*s*_ = 0.001, *p*_*e*_ = 0.004, resulting in a small number of larger clusters. Colours in B,C,D correspond to these parameter settings. (B) Distribution of cluster sizes for 1000 independent simulations for each parameter setting on a logarithmic scale. Distribution of blue does not exceed cluster size 20; (C) Distribution of instantaneous speed of 1000 independent simulations for each parameter setting. Mean speed for orange is lower compared to blue; (D) Log-log plot of the MSD. The onset of the second linear regime (log-log slope approximately equal to 1) is marked with an arrowhead in corresponding colours. This regime starts at smaller *dt* for the orange curve compared to the blue curve, which corresponds to a lowered persistence time compared to blue. Simulations are also available as S3 Video.

Next, we studied whether the differences in adhesion distribution lead to differences in cell motility. First, the speed distribution has a slightly higher mean for the many-small-cluster (blue) setting, and appears more bimodal than the few-large-cluster (orange) setting (Fig 8C). The MSD curves largely overlap on the short term, but on the long term the few-large-cluster (orange) setting shows an earlier start of the final linear regime. The onset of this regime corresponds to the persistence time, which we obtained by fitting the MSD curves with Eq 2 as well as Eq 4. Indeed, the few-large-cluster (orange) setting has about 25% lower persistence time than the many-small-cluster (blue) setting, possibly because it is harder to detach a large adhesion patch than a few small ones. So not only the total adhesion area influences cell motility, but also how that area is distributed over adhesion clusters and where those clusters are located. This further shows that the dynamics of cell-matrix adhesion influence cell motion and can be a key component of the plasticity of cell motility.

## Discussion

Lymphocytes show a large variety of motility behaviours *in vivo* and on substrates. This variety can be partially ascribed to the substrate composition. Another part is caused by the properties of individual cells. To understand how the cell properties and substrate properties give rise to the large variety observed in lymphocytes, we propose a novel extension of the CPM-Act model with dynamic cell-matrix interactions.

In the presented model, cell-matrix adhesions can develop *de novo* in an Act-dependent manner, and adhesion patches can shrink and grow. Furthermore, adhesions can break for a set energy cost. Together with the Act-dependent cell propulsion, this model is sufficient to describe the floating, stepping, and pivoting behaviour observed in B cells on fibronectin [12], by just altering adhesion bond energy cost and *de novo* adhesion formation, as well as the walking and gliding behaviour observed in T cells on ICAM and casein [13], by adjusting *de novo* adhesion formation and adhesion patch growth.

We first showed that increasing the strength of adhesion bonds, *λ*_*adh*_, and increasing the probability by which adhesions form *de novo, p*_*s*_, decreases cell speed and especially cell persistence. For very strong bonds, the cells remain attached to the matrix while trying to move away. Such motility is persistent on a short time scale, but subdiffusive on long-time scales. With the addition of adhesion-dependent cell propulsion strength, a richer behavioural repertoire can be reproduced for low adhesive areas: simulated cells with very low adhesive areas have low diffusion, as their propulsion strength is weakened, whereas cells with slightly higher adhesive areas can show temporary spurts of increase in adhesion area combined with increase in speed. Finally, we showed that the distribution of adhesion clusters affects cell motility. Cells with many small adhesion clusters at the cell front have a higher persistence time than cells that have fewer but larger adhesion clusters located at the centre and the back of the cell, even while total adhesion area is equal.

### Linking cell variability and parameter variability

The experimental studies by Rey-Barroso et al. [12] and Jacobelli et al. [13] showed variability in motility among the cells. Other studies have also described variability among genetically identical *Dictyostelium discoideum* cells in their chemotactic performance [36], and among keratocytes in their shape and speed. Which features of the individual cells underlie such variability in cell motility? By adjusting the adhesion formation rate, adhesion strength and adhesion distribution, we could already capture the different motility modes of B cells on fibronectin and of T cells on casein and ICAM Still, it would be very interesting to be able to link the changes in these parameters to actual differences between cell populations on different substrates or between individual cells on the same substrate.

An interesting starting point for addressing the variability in lymphocyte motility are measurements of the different subsets of differentiated CD4+ T cells. Th1, Th2, and Th17 subsets have been described to harbour distinct motility properties both *in vitro* and *in vivo*, as well as distinct molecular equipment in terms of adhesion and cytoskeleton dynamics [37, 38]. Th1 and Th17 cells show low speed and displacement, have a low expression of integrin *α*_*v*_*β*_3_ and show fluctuating, yet high concentrations of Ca^2+^, whereas Th2 cells have high mobility, high levels of *α*_*v*_*β*_3_ and lower and more constant levels of [Ca^2+^]. Interestingly, these differences have been proposed to support distinct search strategies aligning with the fact that these cell subsets target different types of pathogens. The differences in motility and integrin expression correspond well with the observations in our simulations with adhesion-dependent protrusions, where a larger rate of *de novo* adhesion resulted in higher motility as well as more adhesive surface. As such, our model predicts a larger adhesive area for Th2 cells than for Th1/Th17 cells, which can be verified experimentally. By further measurements such as measuring integrin expression levels among individual cells by flow cytometry, or monitoring size and distribution of adhesions with super-resolution microscopy approaches, we can specify our model parameters and assess the predictive value of our model. In general, our study provides a mechanistic framework to identify which processes lead to differential motility and then address this experimentally.

Conversely, experimental measurements can elucidate the current estimates of protrusion and adhesion energies. With the current parameter settings, protrusion energy exceeded the adhesion energy, whereas our literature-based estimates suggest the reverse should be true (S2 Material, [39–42]). However, the references studies measured adhesion forces in fibroblasts, osteoclasts and CHO cells, which can have adhesive properties distinct from lymphocytes. For instance, a study on adhesion between a T cell and TNF-*α* stimulated HUVEC cells measured a de-adhesion work of the entire T cell in the same order as our estimates for protrusion-associated work of a single lattice site [43]. This suggests that our estimated adhesion energies based on other cell types might be largely overestimated. However, how this cell-cell adhesion energy translates to adhesion energies on substrates is yet unclear, and new experimental measurements on lymphocyte adhesion energies on substrates can further improve our model.

### Motility on multiple time scales

An important feature of our model is the possibility to study cell motility at multiple timescales, especially long-term motility. For long-term behaviour in our model, we mostly observed regular diffusive behaviour or, for more extreme *λ*_*Act*_ and *p*_*s*_ values, subdiffusive behaviour. This corresponds with the pivoting B cells observed by Rey-Barroso et al. [12]. The other extreme, superdiffusive behaviour, has also been observed in mammalian cells. The murine T cells in Harris et al. [4] showed superdiffusive behaviour, but have only been tracked for a relatively short time (*∼*10 min), so their diffusive behaviour on longer time scales is unknown. In comparison, our simulated cells display persistent, and hence superdiffusive, behaviour at time scales up to 400 to 1000 MCS, which is in the order of minutes, yet they display regular diffusive behaviour at larger timescales. This highlights the existence of multiple time scales in cell motility. Interestingly, the Madin-Darby canine kidney cell in Dieterich et al. [32] have also been tracked for longer time (*∼*1000 min) and show three time scales of roughly 0-4 minutes, 4-16 minutes and from 16 minutes onwards, at all of which superdiffusion is observed. Furthermore, cell velocities show long range correlations in time. What causes these long-time correlations and the corresponding long-term superdiffusive behaviour is unclear.

Although our model provides a plausible explanation for the impact of cell adhesion on subdiffusive cell motility, it does not reproduce the observation that B cells move faster and in a Brownian fashion on fibronectin, but slowly and persistently on collagen IV [12] at the intermediate time scale. Based on the observation of higher adhesion areas on fibronectin than on collagen, we could argue that this can be associated with higher *p*_*s*_ and *p*_*e*_ in our model, as these parameters determine adhesion area. Yet, for increasing adhesion areas above the full activation of protrusions, we see a drop in both speed and persistence, whereas for increasing adhesion areas while below the fully activated protrusions, both speed and persistence increase. Overall, in the model, speed and persistence are highly correlated: fast cells show high persistence and slow cells are not persistent. An experimental study showed this correlation to be a universal coupling between cell speed and cell persistence (UCSP), mediated by actin flow [44], as actin flow stabilizes cell polarity. In our model, the actin flow is modelled phenomenologically by the Act model [19], which displays this UCSP as well [20]. So the observation that B lymphocytes can display slow persistent motility on fibronectin, and much faster Brownian cell motility seems to disagree with UCSP. Signalling between the cytoskeleton and the matrix may further contribute to the differences in cell motility between the two types of matrices.

### Interplay between cytoskeleton and substrate

In our current model, there is only an explicit interaction between the adhesions and the Act-extension. Specifically, only the protrusion efficiency is directly influenced by the presence of adhesion. However, whether other aspects of the actin network, such as the actin lifetime or degradation rate, are influenced by adhesions is not taken into account.

Where the Act-extension mainly models the actin-network at the front of the cell, many locomotion-related processes also involve the contractile components of the cytoskeleton, such as myosin-II. Myosin-II contraction pulls the back of the cell towards the front and can increase cell speed [13]. Within the CPM, myosin-II contractility is modelled indirectly through the perimeter constraint and the contact energy between cell and medium. Changing parameter values for both results in altered speeds within our model (S4 Video). Part of this cortical tension is transferred onto the matrix through adhesions [45, 46]. An interesting question is whether the cortical tension is also influenced by the presence of adhesions. Furthermore, myosin-II is suggested to be a polarization cue and to be transferred to the back of the cell by retrograde actin flow and could possible also alter persistence of cell polarization [44]. An interesting direction for future research would be to study how the retrograde flow is influenced by cell-matrix adhesions and how this may affect the UCSP that also depends on the actin flow.

Next, the retraction of the rear end is slowed down by adhesions if they do not detach. Our current model uses greatly simplified adhesion patch detachment: a stochastic process of loss of sites, and energy requiring retractions of adhesion sites. It ignores the following two processes: First, myosin-II, besides contracting the rear end, is also involved in the detachment of adhesion patches in T cells [13, 47], as it increases the forces exerted on the adhesions. Second, adhesion detachment at the rear of the cell is also regulated by scaffolding proteins talin an moesin. Both compete for binding sites between integrin and the cytoskeleton, but have different properties. While talin connects the cytoskeleton to integrins, moesin inactivates integrin, thereby decoupling the adhesion from the cytoskeleton at the rear end [48]. Considering the myosin-II dependent detachment and rear-end specific detachment in a next model can further enhance the understanding of the role adhesion cluster size and distribution in cell motility.

So far, we have addressed model improvements regarding intracellular processes. However, cell-matrix interactions are also determined by integrin and matrix properties, including mechanistic feedback between the integrins and matrix. Hence, both matrix rigidity and the cell’s ability to generate force influence cell shape and cell motion. When it comes to modelling this feedback, different approaches have been used already in phase-field models of cell motility. In Copos et al. [15], adhesions were modelled as mechanosensitive bonds. In Ziebert et al. [16], adhesions ruptured when they exceeded a maximum length. In Shao et al. [49] the probability of adhesion rupture increased with force. In Löber et al. [17], the matrix deformation was also taken into account, leading to non-trivial motion such as bipedal motion. In Kim et al. [50], a 3D matrix was modelled as individual fibres which a cell can push and pull on. This allows filopodia to sense the matrix stiffness locally, which results in durotaxis.

Modelling matrix deformation or displacement of adhesion sites within the CPM is challenging, but a lot of progress has been made recently. Methods to estimate forces within the CPM cell have been developed, either based on cell shape or on the Hamiltonian [51, 52]. The CPM has also been combined with a finite element method to model matrix traction forces with feedback between the CPM and FEM [51, 53]. Explicit descriptions of cell-matrix adhesions were recently introduced into this framework [54] to describe tension-dependent growth of focal adhesions. In our future work, this methodology will allow us to study the mechanisms by which substrate compliance affects lymphocyte motility.

In conclusion, we propose a simple mechanism by which the ECM can affect the characteristics of cell trajectories. To this end, we have introduced a novel CPM model that combines the Act model [19] with dynamic cell-matrix adhesions, generating a large repertoire of cell trajectories (Figure 9)We show that a simple model considering actin-driven protrusion formation in interaction with the dynamic formation and detachment of adhesions to the substrate, suffices to reproduce both persistent random walks as well as short-term persistent but long-term subdiffusive random walks. Thus our simplified model reproduces the motility patterns observed in individual B cells on a fibronectin substrate, such as reduced motility for non-attached cells, walking motility, and pivoting motility due to sustained attachment, as well as the walking and gliding motion of T cells on ICAM or casein substrate. The computational efficiency of the model allowed us to efficiently study both short-term molecular scales as well as the long-term cell behaviour following from it, providing insight into the molecular parameters that explain the plasticity of cell motility due to interaction with substrates. Our study shows that the interplay between adhesion formation, adhesion expansion and adhesion strength may determine the turn-over of the adhesion area which regulates cell speed and persistence.

**Fig 9.**
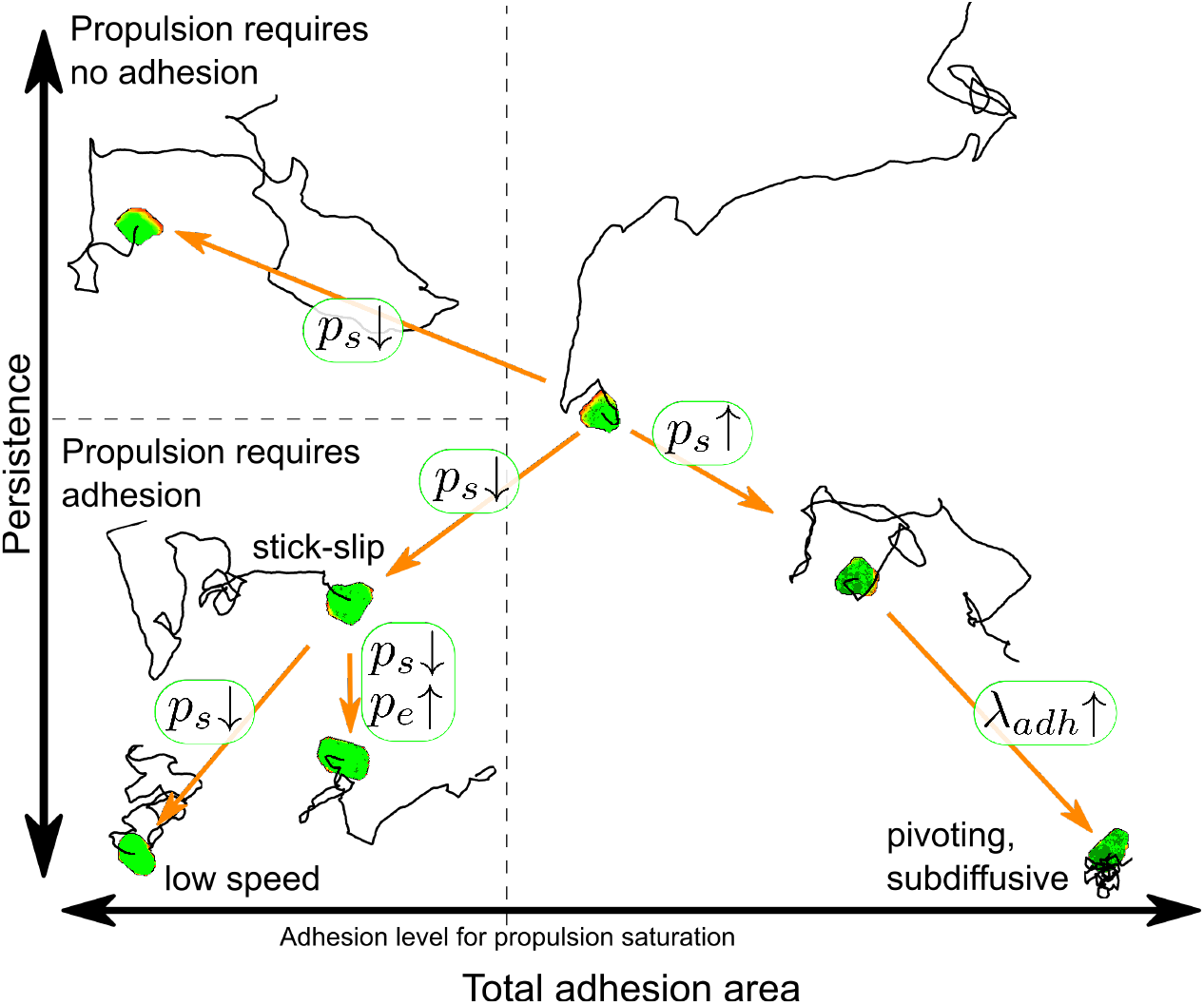
Overview of the motility modes possible in the model and which parameters govern the transitions between them. For each motility mode, a representative cell and its trajectory are plotted in a persistence versus total adhesion area plane.

## Methods

In this work, we model the cells moving on and adhering to flat substrates. The basis of our model is the Cellular Potts model.

### Cellular Potts model

We use the Cellular Potts model (CPM) to represent a cell on a regular, square lattice. Lattice sites 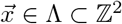 are assigned an identity 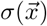, with 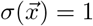 if the site belongs to the cells, and 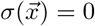 if the cell belongs to the medium. The cell can then be defined as the set 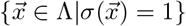. The model mimics cell protrusions and retractions through iterative attempts to extend or retract the cell into one of the neighbouring lattice sites, depending on a Hamiltonian function, *ℋ*, representing passive forces acting on the cell, and a number of active, dissipative processes (e.g., actin protrusion). Together, these represent the balance of forces that drive cell motility. More formally, the algorithm selects a pair of adjacent lattice sites 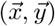, where cells are considered adjacent if they are adjacent orthogonal or diagonal neighbours, i.e.,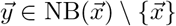 where 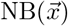 is defined as the set of the eight first and second-order neighbours of 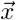.

The Hamiltonian, *ℋ*, describes the balance of passive forces produced acting on the cell in terms of a system energy,

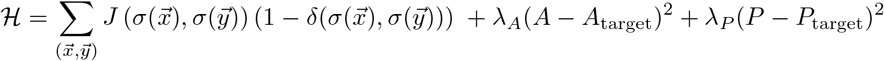

The first term describes the contact energies between cell and medium. The second term describes an area constraint, with *A* and *A*_target_ being the area and target area of the cell and, similarly, the third term describes a perimeter constraint with *P* and *P*_target_ the perimeter and target perimeter of the cell.

The total change in energy is given by the change in the Hamiltonian, Δ*ℋ*due to the attempted update in addition to the energy, Δ*ℋ*_*Act*_ coming from the Act model and Δ*ℋ*_*Adh*_ giving the energies associated with the detachment of cell-ECM substrate adhesions. The total change in energy Δ*ℋ*_total_ = Δ*ℋ*+ Δ*ℋ*_*Act*_ + Δ*ℋ*_*Adh*_ determines the acceptance probability of a copy attempt,

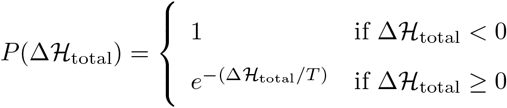

Here, *T* controls the amount of random fluctuations in the system. Higher *T* will allow more thermodynamically unfavourable copy attempts to be accepted. In the Cellular Potts model, time progresses in Monte Carlo step (MCS), which represents a unit of time that allows each lattice site to be updated once on average.

## Cell motility - Act model

Cells move by making protrusions through actin polymerization and form cell extensions like filopodia, pseudopodia, and lamellipodia. Actin polymerization in the CPM has previously been modelled in a phenomenological way in the Act model by Niculescu et al. [19]. This extension adds an extra layer to the CPM, describing the Act values of lattice sites, ranging from 0 to maximum value Max_*Act*_. For lattice site 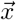 newly added to the cell, 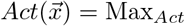. At each MCS, the Act values are decreased by 1 until 0. The term 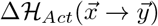 is subtracted from Δ*ℋ*, and can be interpreted as the resulting force from pushing and resistance at the membrane element between 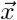 and 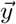. In Δ*ℋ*_*Act*_, the local geometric mean of Act-values of both the expanding and retracting lattice sites are compared and the lattice site with the highest mean is favoured in the following way:

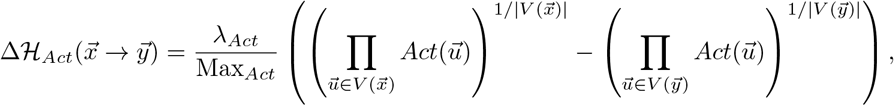

with 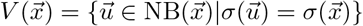 the neighbourhood of lattice site 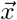 restricted to the same cell, and *λ*_*Act*_ is the weight given to this model component.

Adhesion to the matrix is required to fully translate actin polymerization to cell membrane protrusion [33–35], as it transmits the polymerization force to the matrix. We add feedback between the cell adhesions and actin polymerization, by increasing the force produced by polymerization upon increase in adhesion area. This is only up to a threshold adhesion area, after which the protrusion force remains fully activated. We therefore multiply *λ*_*Act*_ with factor *f* defined as follows,

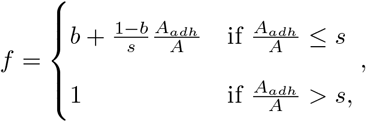

Here, *A* denotes the area of the cell, *A*_*adh*_ the adhesive area of the cell, *b* the value of *f* when there are no adhesions, and *s* the fractional adhesive area at which *f* saturates.

### Cell-substrate adhesions

The adhesions of a cell to the extracellular matrix are modelled as a third layer in the CPM. A lattice site 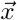 in this layer can either contain no adhesion 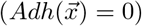, or an adhesion patch 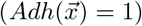. Adhesion dynamics are governed by four processes: *de novo* formation of adhesions, adhesion patch expansion, adhesion patch unbinding, and rupture of adhesion through retraction of a cell. We describe each of these processes below.

#### New adhesion sites

New adhesions form when the cell membrane comes in close enough contact with the extracellular matrix such that integrins can bind to the matrix. This process is dependent on actin polymerization, membrane protrusion and polarized distribution of integrins [24–26]. We model *de novo* adhesion formation through a stochastic process. Each MCS a grid site within a cell can turn from non-adhesion to an adhesion site with probability *p*_*s*_, if the local geometric mean restricted to the cell of the Act layer exceeds the value 0.75 Max_*Act*_, i.e.:

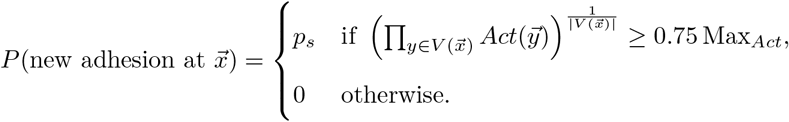

#### Adhesion patch expansion

Once adhesion patches are formed, they can increase in size. Multiple processes underlie this expansion. First, once the cell membrane is attached to the matrix, it fluctuates less, allowing for easier attachment of new integrins [27]. Secondly, the curvature of the cell membrane favours aggregation of integrins [55, 56].

We do not model integrin recruitment and membrane curvature, but choose to model adhesion patch growth phenomenologically. Jacobelli et al. [13] observed that adhesion patches grow radially, with some bias in the direction of the cell front. Hence, we model adhesion patch expansion as an Eden-like growth model [28], known to give roughly circular shapes. While updating the adhesion layer, once a lattice site containing an adhesion is selected to be updated, we also select a random neighbour. If that neighbouring lattice site contains no adhesion, it forms an adhesion with probability *p*_*e*_.

#### Adhesion patch unbinding

Aside from patch expansion, patch unbinding can also occur, either spontaneously [57] or influenced by myosin-II contraction [13]. Following the observation of concentrical patch detachment [13], an adhesion site 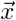 in this model can spontaneously detach with a probability depending on the adhesion status of its neighbours.

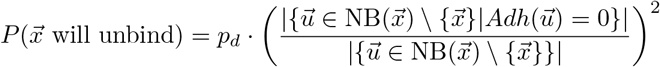

with 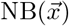 the Moore neighbourhood of 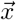. Thus, the higher the number of non-adherent neighbours, the higher the probability that the site loses its adhesion.

#### Adhesion rupture through retraction

Adhesions at the cell rear can also unbind by force. Although integrins are known to show catch-slip bond behaviour [58, 59], we simplify the rupture of an adhesion to a constant amount of energy required to break an adhesion upon cell retraction. This determines the term contributing to 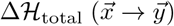:

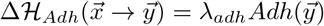

with 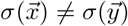 and the cell 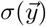 retracting.

### Implementation

A measure of time in the CPM is the Monte Carlo Step (MCS). Within one MCS, the expectation is that the *σ* of each lattice site has been updated once. However, many of the proposed neighbouring lattice site pairs share the same *σ* and will thus not result in a changed model state. Therefore, we use a rejection-free algorithm that only considers attempts between neighbours of different *σ* to speed up simulations [60, 61]. Further, the adhesion layer and Act layer of the model are also updated during and after the *σ*-update. Act-values and adhesion updates regarding the relocation of the cell are executed immediately during the *σ*-update: e.g., for copy attempts that let a cell retract from a lattice site, we do directly update the Act-values and adhesions of that site. After the *σ*-update, we update the adhesion layer asynchronously: we iterate, in random order, over the lattice sites within the cell and execute the processes described in the Cell-substrate adhesions subsection. Lastly, we update the Act-layer: every Act-value is diminished by 1 until 0. These three updates together constitute one MCS. The model has been implemented in Tissue Simulation Toolkit and is available in S1 Material.

### Simulation parameters

During our different simulations, many parameter values were kept constant (Table 3). All simulations were done on a 300 × 300 lattice with periodic boundaries with a single cell. Parameter values that were not constant are shown in Table 1. For the simulations in Fig 2, 3 and 4, *p*_*d*_ = 0.0008, and *p*_*s*_ and *λ*_*adh*_ varied according to the figure legends. For simulations shown in Fig 6 and 7, *p*_*d*_ = 0.001 and again *λ*_*adh*_ varied according to the figure legends. The Act-only simulations in Fig 3 and Fig 7 were run with all adhesion dynamics parameters equal to zero: i.e., *λ*_*adh*_, *p*_*s*_, *p*_*e*_, and *p*_*d*_ were all zero. For all simulations, *λ*_*Act*_ = 240, except for the specific Act-only simulations in Fig 7 with *λ*_*Act*_ = 120. For the simulations in Fig 8, *p*_*e*_ and *p*_*s*_ were varied, see figure legend. The parameters not mentioned in the figure legend are *p*_*d*_ = 0.0004, *λ*_*adh*_ = 60, *b* = 0.5, *s* = 0.12.

**Table 3.** List of parameter values kept constant during all simulations. Values are arbitrary units, unless specified otherwise.

| Parameter | Description | Value |
| --- | --- | --- |
| $T$ | temperature | 30 |
| $A_{\text{target}}$ | target area | 1000 $px^2$ |
| $\lambda_A$ | weight area constraint | 50 |
| $P_{\text{target}}$ | target perimeter | 350 $px$ |
| $\lambda_P$ | weight perimeter constraint | 4 |
| $\lambda_{Act}$ | weight of Act model | 240 |
| $\text{Max}_{Act}$ | Act lifetime | 120 *MCS* |
| $J_{\text{medium,medium}}$ | adhesion energy between medium | 0 |
| $J_{\text{cell,medium}}$ | adhesion energy between cell and medium | 35 |
| Total MCS | simulation duration | 25000 *MCS* |

## Acknowledgments

Martijn de Jong is thanked for his implementation of the rejection-free algorithm of the Cellular Potts model in the Tissue Simulation Toolkit. G.T. gratefully acknowledges the Indian Institute of Science to serve as Infosys visiting professor at the Centre for Ecological Sciences in Bengaluru. L.v.S. would like to thank the French Embassy in the Netherlands for the Bourse d’Excellence Descartes de stage. R.M.H.M. is supported by Nederlandse Organisatie voor Wetenschappelijk Onderzoek grant NWO/ENW-VICI 865.17.004. We also thank SURFsara for the support and computing time in using the Lisa cluster computer.

## Supporting information

**S1 Figure Fürth with translational diffusion fits MSD of model better than Fürth without translational diffusion**. Log-log plot of MSD for the four scenarios in Fig 2, similar to Fig 4, with fits of Eqs 1 and 2. Parameters are: A) *λ*_*adh*_ = 20, *p*_*s*_ = 0.004, B) *λ*_*adh*_ = 100, *p*_*s*_ = 0.004, C) *λ*_*adh*_ = 20, *p*_*s*_ = 0.02, D) *λ*_*adh*_ = 100, *p*_*s*_ = 0.02.

**S2 Figure Persistence times obtained from fitting MSD with Fürth with translational diffusion (Eq 2) against adhesion area for different values of** *p*_*s*_ **and** *λ*_*adh*_. Parameters are the same as in Figure 3, except that *λ*_*adh*_ has been limited to 20, 40, 60 because of bad fitting with Eq 2. For reference, the persistence time of the Act model without the adhesion extension is plotted as the black dot.

**S3 Figure MSD for initial and extended model with fits of Eq 2 and Eq 4** Parameters are the same as in Figure 6, with *p*_*s*_ being varied: (A) *p*_*s*_ = 0.001, (B) *p*_*s*_ = 0.004, and (C) *p*_*s*_ = 0.0025.

**S1 Video Cell speed and persistence drop with increasing values for adhesion formation and adhesion strength**.

**S2 Video Cell speed and persistence drop in the model with feedback from adhesions onto cell protrusion when adhesive areas are small**.

**S3 Video Similar adhesive area sizes lead to different motility when adhesion growth is dominated by the actin-dependent formation of new adhesions versus the growth of existing patches**.

**S4 Video Cell speed is influenced by contractile forces as arise from the perimeter constraint and adhesion energy** *J*_*cell,medium*_. Parameter values are identical to Fig 8, blue except for indicated values.

**S1 Material Model implementation in Tissue Simulation Toolkit S2 Material Estimates of parameter units**

## References

1. Volpe G, Volpe G. The topography of the environment alters the optimal search strategy for active particles. Proceedings of the National Academy of Sciences of the United States of America. 2017;114(43):11350–11355. doi:10.1073/pnas.1711371114.

2. Tejedor V, Voituriez R, Bénichou O. Optimizing persistent random searches. Physical Review Letters. 2012;108(8):088103. doi:10.1103/PhysRevLett.108.088103.

3. Viswanathan GM, Buldyrev SV, Havlin S, Da Luz Mge, Raposo EP, Stanley HE. Optimizing the success of random searches. Nature. 1999;401(6756):911–914. doi:10.1038/44831.

4. Harris TH, Banigan EJ, Christian DA, Konradt C, Tait Wojno ED, Norose K, et al. Generalized Lévy walks and the role of chemokines in migration of effector CD8+ T cells. Nature. 2012;486(7404):545–548. doi:10.1038/nature11098.

5. Bartumeus F, Catalan J, Fulco UL, Lyra ML, Viswanathan GM. Optimizing the Encounter Rate in Biological Interactions: Lévy versus Brownian Strategies. Physical Review Letters. 2002;88(9):4. doi:10.1103/PhysRevLett.88.097901.

6. Miller MJ, Wei SH, Cahalan MD, Parker I. Autonomous T cell trafficking examined in vivo with intravital two-photon microscopy. Proceedings of the National Academy of Sciences of the United States of America. 2003;100(5):2604–9. doi:10.1073/pnas.2628040100.

7. Worbs T, Mempel TR, Bölter J, Von Andrian UH, Förster R. CCR7 ligands stimulate the intranodal motility of T lymphocytes in vivo. Journal of Experimental Medicine. 2007;204(3):489–495. doi:10.1084/jem.20061706.

8. Textor J, Peixoto A, Henrickson SE, Sinn M, von Andrian UH, Westermann J. Defining the quantitative limits of intravital two-photon lymphocyte tracking. Proceedings of the National Academy of Sciences of the United States of America. 2011;108(30):12401–6. doi:10.1073/pnas.1102288108.

9. Espinosa-Carrasco G, Saout CL, Fontanaud P, Michau A, Mollard P, Hernandez J, et al. Integrin β1 optimizes diabetogenic T cell migration and function in the pancreas. Frontiers in Immunology. 2018;9(MAY):1156. doi:10.3389/fimmu.2018.01156.

10. Mrass P, Petravic J, Davenport MP, Weninger W. Cell-autonomous and environmental contributions to the interstitial migration of T cells. Seminars in Immunopathology. 2010;32(3):257–274. doi:10.1007/s00281-010-0212-1.

11. Krummel MF, Bartumeus F, Gérard A. T cell migration, search strategies and mechanisms. Nature Reviews Immunology. 2016;16(3):193–201. doi:10.1038/nri.2015.16.

12. Rey-Barroso J, Calovi DS, Combe M, German Y, Moreau M, Canivet A, et al. Switching between individual and collective motility in B lymphocytes is controlled by cell-matrix adhesion and inter-cellular interactions. Scientific Reports. 2018;8(1):5800. doi:10.1038/s41598-018-24222-4.

13. Jacobelli J, Bennett FC, Pandurangi P, Tooley AJ, Krummel MF. Myosin-IIA and ICAM-1 regulate the interchange between two distinct modes of T cell migration. Journal of immunology (Baltimore, Md: 1950). 2009;182(4):2041–50. doi:10.4049/jimmunol.0803267.

14. Deutsch A, Friedl P, Preziosi L, Theraulaz G. Multi-scale analysis and modelling of collective migration in biological systems. Philosophical Transactions of the Royal Society B: Biological Sciences. 2020;375(1807). doi:10.1098/rstb.2019.0377.

15. Copos CA, Walcott S, del Álamo JC, Bastounis E, Mogilner A, Guy RD. Mechanosensitive Adhesion Explains Stepping Motility in Amoeboid Cells. Biophysical Journal. 2017;112(12):2672–2682. doi:10.1016/j.bpj.2017.04.033.

16. Ziebert F, Aranson IS. Effects of Adhesion Dynamics and Substrate Compliance on the Shape and Motility of Crawling Cells. PLoS ONE. 2013;8(5):e64511. doi:10.1371/journal.pone.0064511.

17. Löber J, Ziebert F, Aranson IS. Modeling crawling cell movement on soft engineered substrates. Soft Matter. 2014;10(9):1365–1373. doi:10.1039/C3SM51597D.

18. Yu G, Feng J, Man H, Levine H. Phenomenological modeling of durotaxis. Physical Review E. 2017;96(1):010402. doi:10.1103/PhysRevE.96.010402.

19. Niculescu I, Textor J, de Boer RJ. Crawling and Gliding: A Computational Model for Shape-Driven Cell Migration. PLoS Computational Biology. 2015;11(10):e1004280. doi:10.1371/journal.pcbi.1004280.

20. Wortel I, Niculescu I, Kolijn M, Gov N, de Boer R, Textor J. Both cell-intrinsic and environmental factors constrain speed and persistence in T cell migration. bioRxiv. 2018; p. 338459. doi:10.1101/338459.

21. Glazier JA, Graner F. Simulation of the differential adhesion driven rearrangement of biological cells. Physical Review E. 1993;47(3):2128–2154. doi:10.1103/PhysRevE.47.2128.

22. Alber MS, Kiskowski MA, Glazier JA, Jiang Y. On Cellular Automaton Approaches to Modeling Biological Cells. In: Mathematical Systems Theory in Biology, Communications, Computation, and Finance. New York, NY: Springer New York; 2003. p. 1–39.

23. Houmadi R, Guipouy D, Rey-Barroso J, Vasconcelos Z, Cornet J, Manghi M, et al. The Wiskott-Aldrich Syndrome Protein Contributes to the Assembly of the LFA-1 Nanocluster Belt at the Lytic Synapse. Cell Reports. 2018;22(4):979–991. doi:10.1016/j.celrep.2017.12.088.

24. DeMali KA, Barlow CA, Burridge K. Recruitment of the Arp2/3 complex to vinculin: coupling membrane protrusion to matrix adhesion. The Journal of cell biology. 2002;159(5):881–91. doi:10.1083/jcb.200206043.

25. Giannone G, Dubin-Thaler BJ, Rossier O, Cai Y, Chaga O, Jiang G, et al. Lamellipodial Actin Mechanically Links Myosin Activity with Adhesion-Site Formation. Cell. 2007;128(3):561–575. doi:10.1016/J.CELL.2006.12.039.

26. Stanley P, Smith A, McDowall A, Nicol A, Zicha D, Hogg N. Intermediate-affinity LFA-1 binds α-actinin-1 to control migration at the leading edge of the T cell. EMBO Journal. 2008;27(1):62–75. doi:10.1038/sj.emboj.7601959.

27. Smith AS, Seifert U. Effective adhesion strength of specifically bound vesicles. Physical Review E. 2005;71(6):061902. doi:10.1103/PhysRevE.71.061902.

28. Eden M. A two-dimensional growth process. Dynamics of fractal surfaces. 1961;4:223–239.

29. Fürth R. Die Brownsche Bewegung bei Berücksichtigung einer Persistenz der Bewegungsrichtung. Mit Anwendungen auf die Bewegung lebender Infusorien. Zeitschrift für Physik. 1920;2(3):244–256. doi:10.1007/BF01328731.

30. Selmeczi D, Mosler S, Hagedorn PH, Larsen NB, Flyvbjerg H. Cell motility as persistent random motion: Theories from experiments. Biophysical Journal. 2005;89(2):912–931. doi:10.1529/biophysj.105.061150.

31. Zeitz M, Wolff K, Stark H. Active Brownian particles moving in a random Lorentz gas. The European Physical Journal E. 2017;40(2):23. doi:10.1140/epje/i2017-11510-0.

32. Dieterich P, Klages R, Preuss R, Schwab A. Anomalous dynamics of cell migration. Proceedings of the National Academy of Sciences of the United States of America. 2008;105(2):459–63. doi:10.1073/pnas.0707603105.

33. Chan CE, Odde DJ. Traction dynamics of filopodia on compliant substrates. Science. 2008;322(5908):1687–1691. doi:10.1126/science.1163595.

34. Hu K, Ji L, Applegate KT, Danuser G, Waterman-Storer CM. Differential transmission of actin motion within focal adhesions. Science. 2007;315(5808):111–115. doi:10.1126/science.1135085.

35. Jurado C, Haserick JR, Lee J. Slipping or Gripping? Fluorescent Speckle Microscopy in Fish Keratocytes Reveals Two Different Mechanisms for Generating a Retrograde Flow of Actin. Molecular Biology of the Cell. 2005;16(2):507–518. doi:10.1091/mbc.e04-10-0860.

36. Samadani A, Mettetal J, Van Oudenaarden A. Cellular asymmetry and individuality in directional sensing. Proceedings of the National Academy of Sciences of the United States of America. 2006;103(31):11549–11554. doi:10.1073/pnas.0601909103.

37. Weber KS, Miller MJ, Allen PM. Th17 Cells Exhibit a Distinct Calcium Profile from Th1 and Th2 Cells and Have Th1-Like Motility and NF-AT Nuclear Localization. The Journal of Immunology. 2008;180(3):1442–1450. doi:10.4049/jimmunol.180.3.1442.

38. Gaylo-Moynihan A, Prizant H, Popović M, Fernandes NRJ, Anderson CS, Chiou KK, et al. Programming of Distinct Chemokine-Dependent and -Independent Search Strategies for Th1 and Th2 Cells Optimizes Function at Inflamed Sites. Immunity. 2019;51(2):298–309. doi:10.1016/j.immuni.2019.06.026.

39. Palecek SP, Loftust JC, Ginsberg MH, Lauffenburger DA, Horwitz AF. Integrin-ligand binding properties govern cell migration speed through cell-substratum adhesiveness. Nature. 1997;385(6616):537–540. doi:10.1038/385537a0.

40. Gallant ND, Michael KE, García AJ. Cell adhesion strengthening: Contributions of adhesive area, integrin binding, and focal adhesion assembly. Molecular Biology of the Cell. 2005;16(9):4329–4340. doi:10.1091/mbc.E05-02-0170.

41. Taubenberger A, Cisneros DA, Friedrichs J, Puech PH, Muller DJ, Franz CM. Revealing Early Steps of α ¡sub¿2¡/sub¿ β ¡sub¿1¡/sub¿ Integrin-mediated Adhesion to Collagen Type I by Using Single-Cell Force Spectroscopy. Molecular Biology of the Cell. 2007;18(5):1634–1644. doi:10.1091/mbc.e06-09-0777.

42. Lehenkari PP, Horton MA. Single integrin molecule adhesion forces in intact cells measured by atomic force microscopy. Biochemical and Biophysical Research Communications. 1999;259(3):645–650. doi:10.1006/bbrc.1999.0827.

43. Zhang X, Wojcikiewicz EP, Moy VT. Dynamic adhesion of T lymphocytes to endothelial cells revealed by atomic force microscopy. Experimental Biology and Medicine. 2006;231(8):1306–1312. doi:10.1177/153537020623100804.

44. Maiuri P, Rupprecht JF, Wieser S, Ruprecht V, Bénichou O, Carpi N, et al. Actin flows mediate a universal coupling between cell speed and cell persistence. Cell. 2015;161(2):374–386. doi:10.1016/j.cell.2015.01.056.

45. Álvarez-González B, Meili R, Bastounis E, Firtel RA, Lasheras JC, Del Álamo JC. Three-dimensional balance of cortical tension and axial contractility enables fast amoeboid migration. Biophysical Journal. 2015;108(4):821–832. doi:10.1016/j.bpj.2014.11.3478.

46. Schiller HB, Hermann MR, Polleux J, Vignaud T, Zanivan S, Friedel CC, et al. β 1 - And α v -class integrins cooperate to regulate myosin II during rigidity sensing of fibronectin-based microenvironments. Nature Cell Biology. 2013;15(6):625–636. doi:10.1038/ncb2747.

47. Jacobelli J, Friedman RS, Conti MA, Lennon-Dumenil AM, Piel M, Sorensen CM, et al. Confinement-optimized three-dimensional T cell amoeboid motility is modulated via myosin IIA-regulated adhesions. Nature Immunology. 2010;11(10):953–961. doi:10.1038/ni.1936.

48. Vitorino P, Yeung S, Crow A, Bakke J, Smyczek T, West K, et al. MAP4K4 regulates integrin-FERM binding to control endothelial cell motility. Nature. 2015;519(7544):425–430. doi:10.1038/nature14323.

49. Shao D, Levine H, Rappel WJ. Coupling actin flow, adhesion, and morphology in a computational cell motility model. Proceedings of the National Academy of Sciences of the United States of America. 2012;109(18):6851–6856. doi:10.1073/pnas.1203252109.

50. Kim MC, Silberberg YR, Abeyaratne R, Kamm RD, Asada HH. Computational modeling of three-dimensional ECMrigidity sensing to guide directed cell migration. Proceedings of the National Academy of Sciences of the United States of America. 2018;115(3):E390–E399. doi:10.1073/pnas.1717230115.

51. van Oers RFM, Rens EG, LaValley DJ, Reinhart-King CA, Merks RMH. Mechanical Cell-Matrix Feedback Explains Pairwise and Collective Endothelial Cell Behavior In Vitro. PLoS Computational Biology. 2014;10(8):e1003774. doi:10.1371/journal.pcbi.1003774.

52. Rens EG, Edelstein-Keshet L. From energy to cellular forces in the Cellular Potts Model: An algorithmic approach. PLoS Computational Biology. 2019;15(12):e1007459. doi:10.1371/journal.pcbi.1007459.

53. Rens EG, Merks RMH. Cell Contractility Facilitates Alignment of Cells and Tissues to Static Uniaxial Stretch. Biophysical Journal. 2017;112(4):755–766. doi:10.1016/j.bpj.2016.12.012.

54. Rens EG, Merks RMH. Cell Shape and Durotaxis Explained from Cell-Extracellular Matrix Forces and Focal Adhesion Dynamics. iScience. 2020;23(9):101488. doi:10.1016/j.isci.2020.101488.

55. Bruinsma R, Goulian M, Pincus P. Self-assembly of membrane junctions. Biophysical Journal. 1994;67(2):746. doi:10.1016/S0006-3495(94)80535-1.

56. Sackmann E, Bruinsma RF. Cell Adhesion as Wetting Transition? ChemPhysChem. 2002;3(3):262. doi:10.1002/1439-7641(20020315)3:3¡262::AID-CPHC262¿3.0.CO;2-U.

57. Lenne PF, Nicolas A. Physics puzzles on membrane domains posed by cell biology. Soft Matter. 2009;5(15):2841. doi:10.1039/b822956b.

58. Merkel R, Nassoy P, Leung A, Ritchie K, Evans E. Energy landscapes of receptor–ligand bonds explored with dynamic force spectroscopy. Nature. 1999;397(6714):50–53. doi:10.1038/16219.

59. Marshall BT, Long M, Piper JW, Yago T, McEver RP, Zhu C. Direct observation of catch bonds involving cell-adhesion molecules. Nature. 2003;423(6936):190–193. doi:10.1038/nature01605.

60. Lee KC. Rejection-free Monte Carlo technique. Journal of Physics A: Mathematical and General. 1995;28(17):4835–4842. doi:10.1088/0305-4470/28/17/016.

61. Starruß J, De Back W, Brusch L, Deutsch A. Morpheus: A user-friendly modeling environment for multiscale and multicellular systems biology. Bioinformatics. 2014;30(9):1331–1332. doi:10.1093/bioinformatics/btt772.

